# Long-read sequencing identifies aberrant fragmentation patterns linked to elevated cell-free DNA levels in cancer

**DOI:** 10.1101/2024.05.02.592182

**Authors:** Benjamin P. Berman, Sarah A. Erdman, Christina Wheeler, Justin Cayford, Jean-Valery Turatsinze, Maria Ouzounova, Marie Piecyk, Marielle Herzog, Léa Payen-Gay, Thomas Walter, Theresa K. Kelly

## Abstract

Circulating cell-free DNA (cfDNA) carries fragmentation patterns that serve as biomarkers of cancer, but standard sequencing approaches miss large portions of the fragment length spectrum. In addition to altered fragmentation patterns, cancer patients often have elevated levels of cfDNA, but the underlying mechanisms are not well understood. To address both questions, we analyzed cancer cases with elevated cfDNA levels using Oxford Nanopore (ONT) sequencing. Long-read ONT sequencing captures the full spectrum of cfDNA fragment lengths and enables cell type inference based on DNA methylation markers. One cohort included cases from several cancer types with elevated cfDNA levels, and a second consisted of patients from a single neuroendocrine cancer study. In each cohort, cases with the highest cfDNA levels showed either hypofragmentation (excess fragments of 1–4 kb) or hyperfragmentation (excess fragments <145 bp). Hypofragmentation reflected blood cell DNA released during delayed sample processing, bearing DNASE1L3-associated hallmarks, while in one cohort we also observed ultra-long fragments (>7.5 kb) lacking these hallmarks and consistent with plasma lysis. By contrast, hyperfragmented samples often had elevated levels of both cancer- and blood-derived DNA, indicating an inflammatory or other system process rather than cancer-specific origin. These findings clarify the distinction between biological and artifactual fragmentation, expand our understanding of cfDNA biology, and highlight long-read sequencing as a powerful tool for biomarker discovery.

## Introduction

Circulating cell-free DNA (cfDNA) is a biomarker for detection and monitoring of cell turnover and death during normal physiological processes and disease [1]. Cancer cells can release cfDNA, and this circulating tumor DNA (ctDNA) can be detected for screening, diagnosis, treatment selection, prognosis, and monitoring [2,3]. Cancer patients often have elevated cfDNA levels, resulting from increases in both cancer- and immune cell–derived cfDNA [4]. Sequencing features used for early cancer detection include mutations and copy number alterations, as well as cancer-specific DNA modifications such as DNA methylation, and changes to the typical cfDNA fragmentation patterns [5–7]. cfDNA fragments typically remain bound to nucleosomes in the blood, allowing some of these changes to be detected by immunological assays or sequencing that targets histone proteins and their post-translational modifications [8–11].

Alterations to the typical fragmentation patterns of cfDNA have been proposed as a cancer marker, but the source of these patterns is not well understood. cfDNA is fragmented by both cellular and circulating endonucleases [12,13]. These fragments often span a single nucleosome, with a characteristic length (∼167 bp) and preference for specific cut sequences (i.e. “fragment end motifs”) [14]. Global alterations in both length [15–18] and end motifs [19,20] have been linked to cancer, but the origin of these changes is poorly understood [5]. In parallel, many fragmentation hotspots are defined by cell type-specific positioning of nucleosomes, and these can be used to detect the presence and cell type of origin of ctDNA [20,21].

Our understanding of these cancer-associated cfDNA fragmentation changes comes primarily from short-read sequencing, which can only resolve relatively small fragments less than 500 bp. Recently, Oxford Nanopore Technologies (ONT) and Pacific Biosciences have been used to sequence longer cfDNA fragments [22–27], which have been observed in fetal DNA, pre-eclampsia, and cancer [23,26,28]. In this study, we use ONT sequencing to survey diverse cancer types and characterize the range of fragmentomic changes, and how they relate to elevated cfDNA and nucleosome levels.

## Results

### Selection of stage III/IV cancer cases with elevated circulating DNA and nucleosomes

In order to select a diverse set of cancer cases for ONT sequencing, we first screened a set of 236 stage III/IV cancer plasma samples from 15 cancer types, as well as 52 plasma samples collected from healthy donors (“Screening set”). We quantified plasma DNA concentration using Qubit, and H3.1 nucleosome concentration using the Nu.Q^®^ H3.1 chemiluminescence immunoassay. H3.1 is a pan-nucleosome marker, and H3.1 levels were highly correlated with total DNA with an R^2^ of 0.73 (Figure 1A). For both cfDNA (Figure 1B) and H3.1 (Figure 1C), most cancer cases were within the normal range, but there were a significant number of cases that were highly elevated. While there were not overall differences in cfDNA or H3.1 levels between Stage III and Stage IV cases (Supplementary Figure 1A), some cancer types did tend to have more elevated cases than others (Figure 1D-E). For instance, about half of Ovarian cancer cases were elevated, whereas very few Head and Neck or Prostate cancers were.

**Figure 1:**
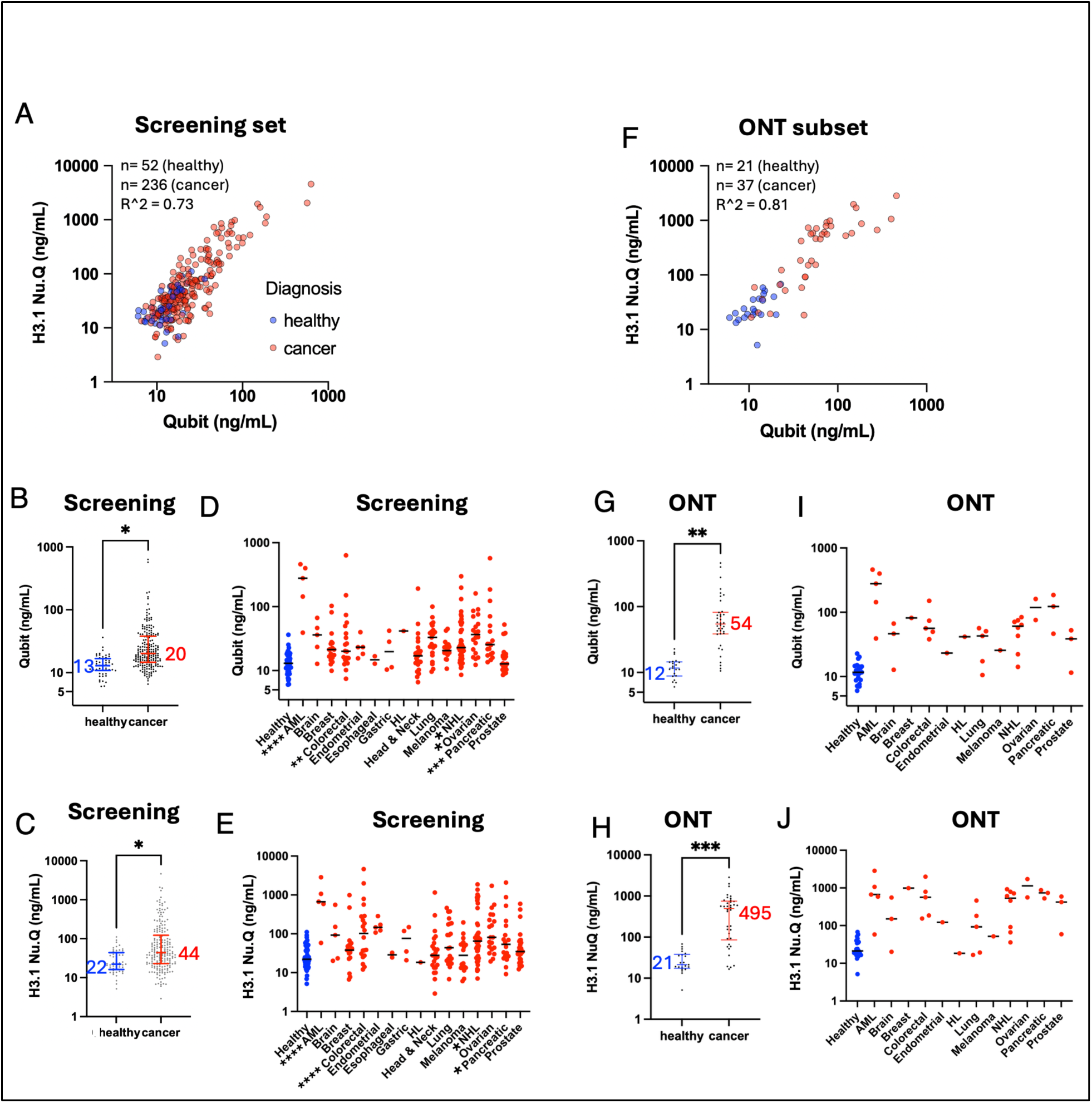
Selection of stage III/IV cancer cases with elevated circulating DNA and nucleosomes. (A-E) A screening set of 52 healthy cases and 236 Stage III-IV cancer cases (“Screening set”). (A) DNA concentration (Qubit) correlated with H3.1 nucleosome concentrations (Nu.Q^®^ H3.1 chemiluminescence immunoassay). (B) DNA concentration for healthy vs. cancer, with median values indicated. (C) H3.1 levels for healthy vs. cancer, with median values indicates. (D) DNA concentration for healthy vs. each cancer type. (E) H3.1 levels for healthy vs. each cancer type. (F-J) Same plots for a subset of 21 healthy samples and 37 cancer samples chosen for Nanopore sequencing (“ONT set”). Error bars show median with interquartile range. P-values in B-E, G, and H were calculated using a two-tailed unpaired t-test (in D-E, any category without an indicator is non-significant). * Signifies p<0.05, ** p<0.01, *** p<0.001.

From this initial screening set, we picked a subset of cancer samples for Oxford Nanopore sequencing (“ONT subset”) that represented the high end of the cfDNA/H3.1 spectrum (mostly with cfDNA above 20 ng/mL and H3.1 above 100 ng/mL, Figure 1F-H). We attempted to represent the full range of cancer types available in our screening set, although some cancer types had very few samples with sufficient DNA for sequencing (Figure 1I-J).

### ONT sequencing detects cancer DNA by copy number alterations (CNA) and methylation cell of origin (COO)

Using ONT MinION 9.4.1 chemistry, we performed whole-genome sequencing [24] with 1-16 million reads per sample (median=4.7 million) (Supplementary Figure 1B). We used ichorCNA [29] to identify Copy Number Alterations (CNAs) and estimate the proportion of tumor DNA in each sample. Using the default parameters, ichorCNA detected cancer DNA in 16 of the 37 cancer samples (Figure 2A). All 21 samples from healthy individuals had an estimated tumor fraction of 0, indicating no false positive CNA calls. While the majority of these cancer samples had very high DNA and nucleosome concentrations, the tumor fraction was not above 0.55 in any case, indicating a major contribution from blood cell DNA and consistent with earlier findings [4].

**Figure 2:**
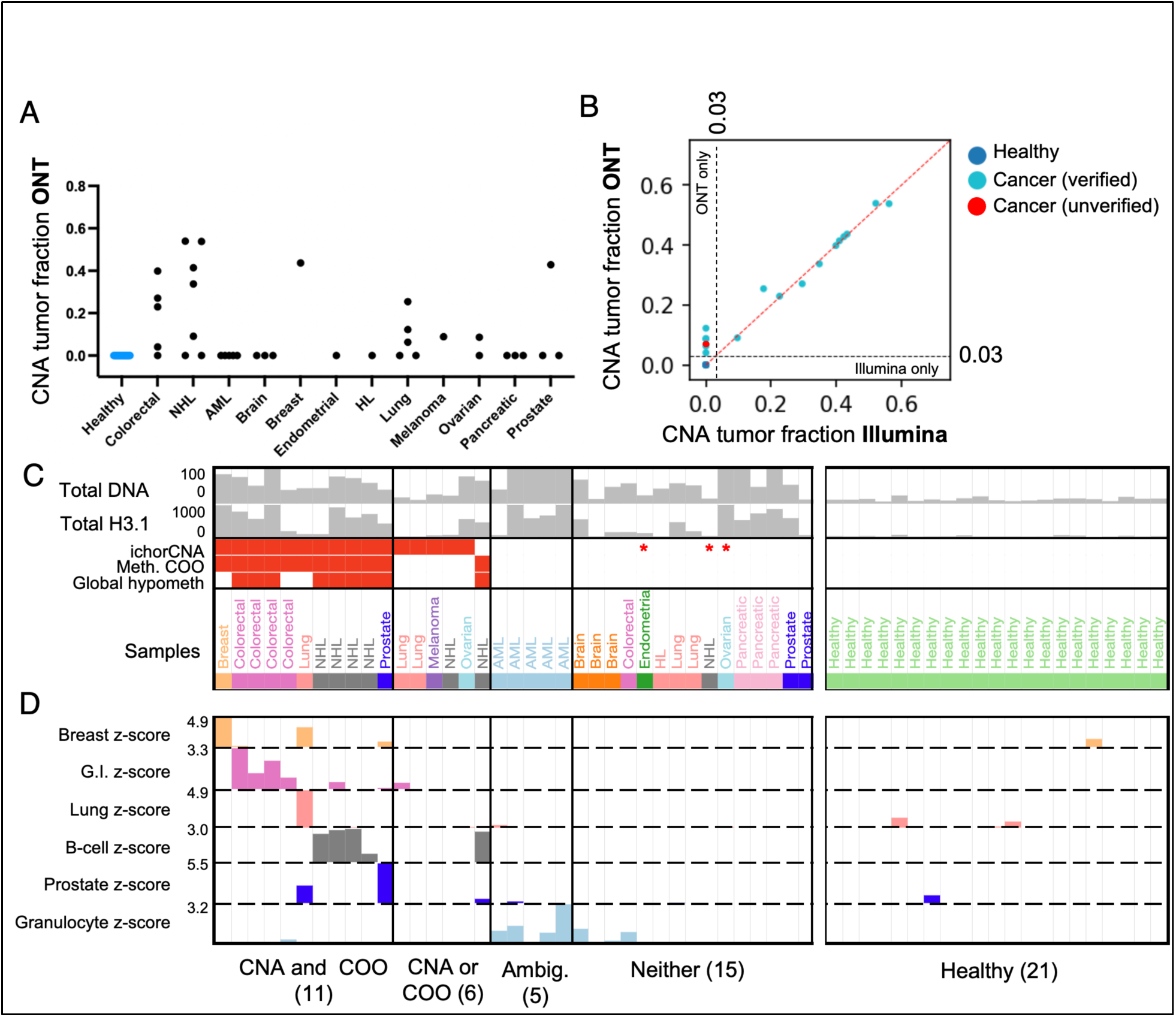
ONT sequencing detects cancer DNA by copy number alterations (CNA) and methylation cell of origin (COO), (A) ichorCNA tumor fractions for 37 samples sequenced using Oxford Nanopore Technologies (ONT). (B) Comparison of ichorCNA-estimated tumor fractions derived from Illumina sequencing (x axis) and ONT (y axis), using default parameters. Dotted line shows the identity line (y=x). Of 5 that were detectable based on ONT but undetectable based on Illumina, all but one could be detected by Illumina with alternative settings (verified), and one was not detectable under any other settings (unverified). (C) Comparison of ONT ichorCNA cancer detection and DNA methylation-based cancer detection. “ichorCNA” indicates whether CNAs were detected by ichorCNA, and Methylation Cell of Origin (“Meth. COO”) indicates that the correct cancer cell of origin had the highest normalized cell type proportion, as shown below for the six relevant cell types. “Global hypometh” indicates that the sample had lower genome-wide methylation than that of healthy samples. Three samples with CNA identified by ichorCNA only with sensitive parameters are shown with asterisks. (D) Normalized cell type fractions from COO analysis are shown for relevant cell types. Values are z-score normalized relative to the healthy samples. Z-score tracks have a lower limit of z=0, and upper limit is indicated in the y axis labels.

We produced matched Illumina sequencing libraries for all healthy samples and 31 of the 37 cancer samples and performed Illumina whole-genome sequencing with 5-50 million read pairs per sample (median 15.8 million read pairs, Supplementary Figure 1C). We ran ichorCNA using default parameters on these matched samples, finding strong concordance in estimated tumor fraction for most samples (Figure 1B). 5 of the 37 cancer samples were called as undetectable in the Illumina data and between 0.06 to 0.12 tumor fraction in the ONT data (“ONT only” samples, Figure 2B and Supplementary Figure 2A). To investigate these further, we constructed a “Panel of Normals” (PoN) for each sequencing platform based the same set of healthy donor samples and ran with ichorCNA settings recommended for low tumor fraction samples [30]. Using these “sensitive” settings, four of the five “ONT only” samples from the original analysis could be recovered from the Illumina data (labeled as “verified” in Figure 2B and shown in detail in Supplementary Figure 2B). The one remaining “ONT only” sample could not be detected under any Illumina settings and may be a false positive call (labeled as “unverified” in Figure 2B). While low tumor fraction samples are often challenging for ichorCNA, ONT provided highly concordant tumor fraction estimates whether or not ichorCNA was run using a Panel of Normals and sensitive settings (Supplementary Figure 2C-E), indicating that ONT WGS might be more robust to sequencing biases.

Using methylation status called directly ONT reads and a methylation atlas of normal cell types, we performed cell of origin (COO) deconvolution to infer the cell type composition of each sample [24,31,32]. Visualizing cell type fractions as z-scores relative to their levels across healthy samples illustrates that the correct cancer cell type was identified for 11 of the 16 CNA-positive samples, and 1 CNA-negative sample (Figure 2C-D). Raw unnormalized cell type fractions confirm the correct cancer cell types (Supplementary Figure 3A). Cell of origin could not be detected for any of the three samples that were detected with ichorCNA sensitive settings but not with the ichorCNA default settings (starred samples in Figure 2C-D). The five Acute Myeloid Leukemia samples had no detectable CNA but did have elevated granulocyte-derived DNA, presumably from cancer cells although this analysis cannot distinguish them from normal granulocytes. The inferred cancer cell of origin fraction was never over 0.5 (Supplementary Figure 3A), reinforcing the idea that blood cells are a major contribution to elevated total cfDNA levels (as quantified directly in Supplementary Figure 3B). Furthermore, 15 of the 37 cancer samples had no detectable ctDNA either by standard ichorCNA analysis or by methylation cell of origin (labeled as “Neither” in Figure 2C), indicating very low cancer fractions despite high cfDNA and H3.1 nucleosome levels in many of these samples. Taken together, these results indicate that many cancers have significant elevation of cfDNA derived from blood cell types.

### Identification of ultra-long sequencing reads likely originating from lysis of contaminating blood cells

Long fragments in cfDNA preps may originate from genomic DNA contamination released during sample processing, especially from cell lysis during freezing and thawing of samples [33,34]. We used frozen plasma samples and performed a post-thaw high-speed spin to reduce the number of long fragments that may originate from contaminating blood cells (Figure 3A). Despite this, three healthy samples had exceptionally high fractions of ultra-long fragments (>7.5 kb) even after the high-speed spin (Figure 3B). In order to understand the origin of these fragments, we compared an aliquot of each of these samples before the high-speed spin step (the “unspun” aliquot). The three outlier samples were highly elevated in cfDNA and H3.1 levels before the spin, whereas they became nearly indistinguishable after the spin (Figure 3C-D). The pre-spin samples also had significantly more ultra-long fragments (>7.5kb) in the ONT sequencing than the post-spin samples (example shown in Figure 3E). As expected, Illumina reads were too short to identify these very long fragments (Figure 3F). In order to be consistent with earlier studies of long cfDNA fragments [26,27], we quantified the fraction of DNA fragments greater than 500, 1000, or 7,500 bp (Figure 3G-I). In each case, the high-speed spin decreased the fraction of long fragments significantly, although this was much more dramatic for fragments >1000bp.

**Figure 3:**
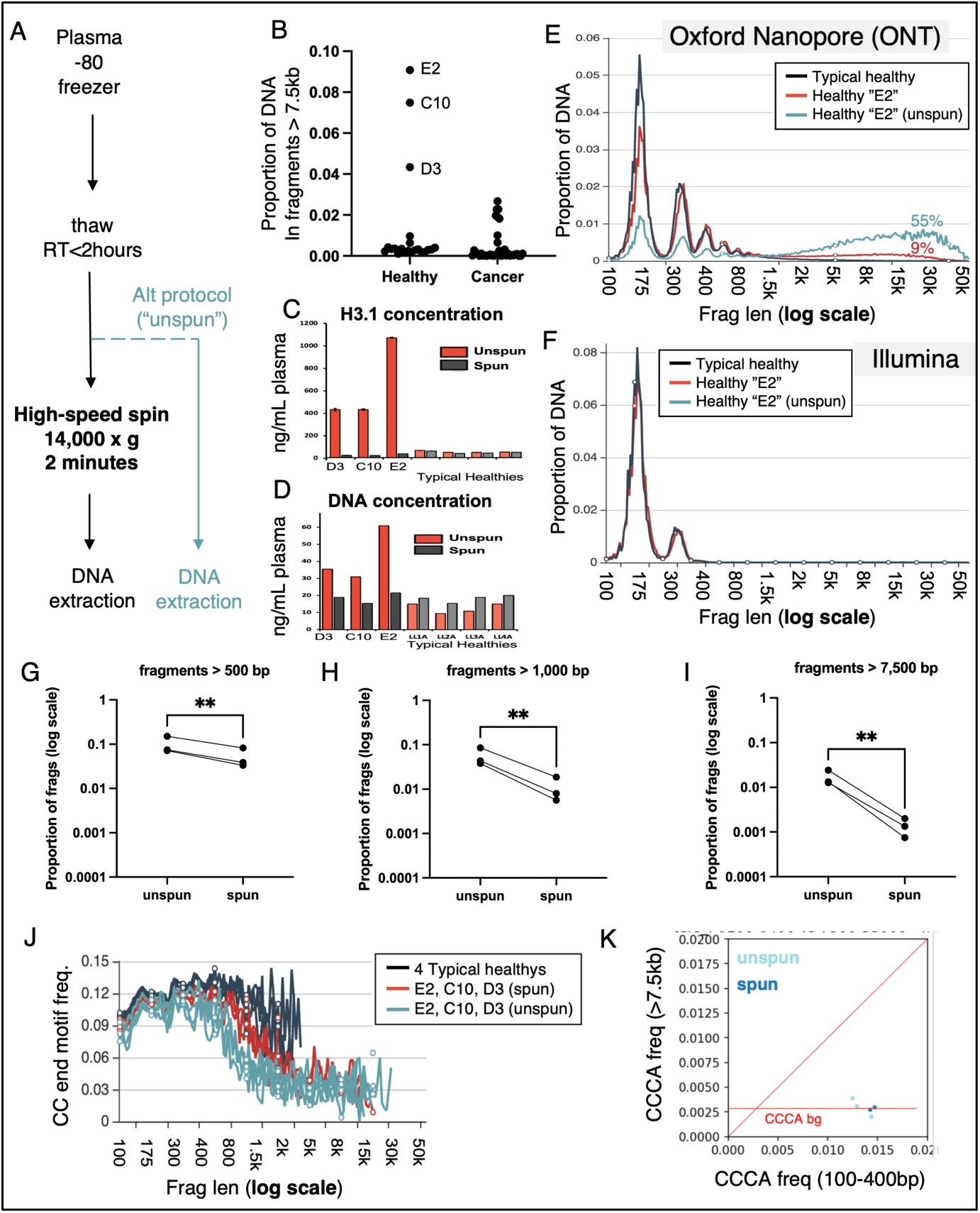
Identification of ultra-long sequencing reads likely originating from lysis of contaminating blood cells . (A) Typical “spun” plasma processing workflow and alternative “unspun” workflow. (B) All ONT samples, showing proportion of DNA in fragments greater than 7.5kb in length. (C) H3.1 concentration and (D) cfDNA concentration for 3 outlier healthy samples and 4 other healthy samples. (E) Proportion of DNA as a function of fragment length, for ONT sequencing of a typical healthy sample (black) and the “E2” healthy sample processed with the spun (red) and unspun (blue) workflows. (F) Proportion of DNA as a function of fragment length, for Illumina sequencing of the typical healthy sample (black) and the “E2” healthy sample processed with the spun (red) workflow. (G-I) Proportion of fragments greater than 500bp (G), 1,000 bp (H) and 7,500 bp (I). (J) Proportion of fragment ends starting with CC in ONT sequencing, as a function of fragment length, for 4 typical healthy volunteers (black) and the three outlier healthy volunteers processed with the spun workflow (red) and the unspun workflow (blue). (K) The proportion of fragment ends starting with CCCA in ONT sequencing, in typical length fragments (100-400bp, x axis) vs. fragments greater than 7.5kb (y axis). In (G-I) ** indicates p<0.01 by two-tailed ratio paired t-test (p values of 0.005, 0.004, and 0.005, respectively).

cfDNA fragments from healthy individuals disproportionally start and end with the nucleotide motif CCCA and others beginning with “CC”, which is attributed to DNASE1L3 activity in circulation [19,35]. To assess whether the ultra-long fragments in our samples were the product of this DNASE1L3-associated fragmentation, we analyzed CC end motif frequencies in the three outlier samples along with 4 typical healthy samples (Figure 3J). Typical healthy volunteers had very few fragments over 1kb to analyze, but those that were present typically had high CC end motif levels (Figure 3J, black lines). In contrast, both the pre- and post- spin samples for the three outlier samples had high CC end motif levels up to about 500bp but decreased precipitously in fragments over 1kb (Figure 3J, blue and red lines). We confirmed that the most specific DNASE1L3-associated motif, CCCA, had high levels in short fragments but very low levels in ultra-long fragments (Figure 3K). These fragments likely represent DNA released from lysed cells in and remaining unfragmented, as observed in earlier studies [33,34].

### Unsupervised clustering by fragment length identifies both hypofragmented and hyperfragmented cancer samples

In order to assess fragment length differences systematically, we performed Principal Component Analysis (PCA) based on fragment length distributions to get an unsupervised clustering of all samples. We used the first 3 principal components (PC1-3, Supplementary Figure 4A-C) to divide the samples into 3 groups of cancers and 3 groups of healthy samples (Figure 4A), which we describe in detail below.

**Figure 4.**
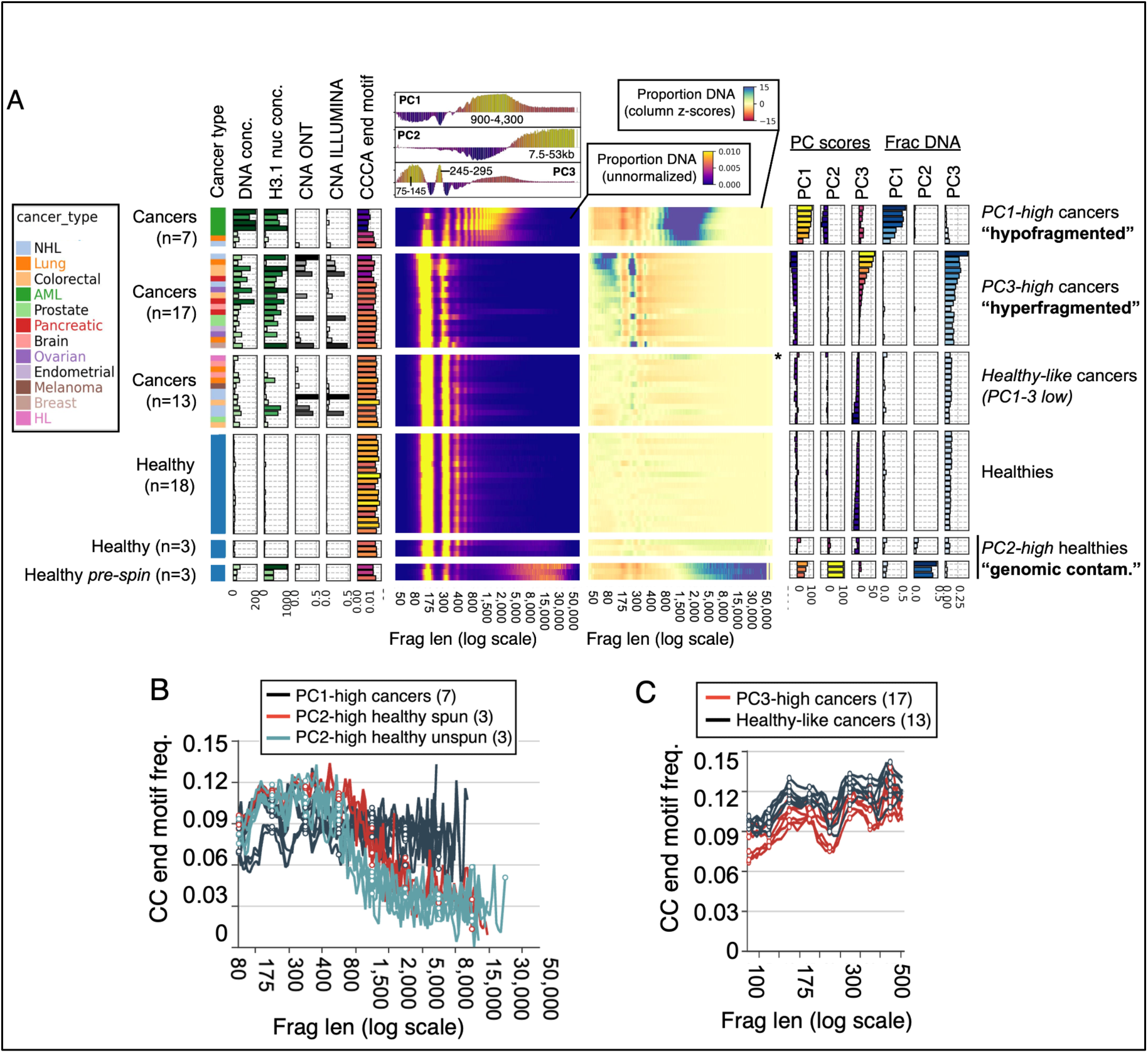
Unsupervised clustering by fragment length identifies both hypofragmented and hyperfragmented cancer samples. (A) All samples ordered by first three Principal Components (PCs). The first 3 sample groups are cancer samples with the cancer type indicated by color, and the second 3 groups are healthy samples. Bar plots on the left show total cfDNA concentration (ng/mL), total plasma H3.1 nucleosome concentration (ng/mL), tumor DNA fraction from ichorCNA from ONT sequencing (“CNA ONT”) and Illumina sequencing (“CNA ILLUMINA”), and the fraction of ONT fragments ending with the motif CCCA. In all bar plots, taller bars are also indicated with darker colors. The heatmap to the right of these bar plots shows the proportion of DNA sequenced in different fragment length bins, on a log scale from 50 bp to 50,000 bp. Above this heatmap are the PC loadings for each of the first 3 PCs. The heatmap to the right shows the same data, but each fragment length bin/column is z-score normalized according to the mean and standard deviation across all healthy samples (not including “pre-spin” samples). The bar plots on the right show the eigenvalues for the 3 principal components (“PC scores”), and the percentage of sequenced DNA in the ranges defined the by those PCs (“Frac DNA”). One sample with a sub-threshold but positive PC1 value is marked with an asterisk. (B) CC end motif frequencies as a function of fragment length, for the PC1 high and PC2 high samples. (C) CC end motif frequencies as a function of fragment length, for the 6 cancer samples with the highest PC3 scores, and the 6 cancer samples with the lowest.

PC1 was most enriched in fragments length 900-4,300bp (Supplementary Figure 4C), and the first sample group (Figure 4A, “PC1-high cancers”) had severely “hypofragmented” DNA in this length range. The cancers in this group had between 20-66% of all DNA in fragments 900-4300bp (Figure 4A, “Frac DNA in PC1”). Interestingly, the PC1-high group included all five Acute Myeloid Leukemias in our study, as well as one Lung cancer and one Non-Hodgkins Lymphoma (NHL) case. An additional Hodgkin’s Lymphoma (starred) was weakly positive for the PC1 signature.

PC2 was most enriched in fragments length >7.5kb, coinciding with the ultra-long fragments of our earlier analysis. PC2-high samples consisted of the spun and unspun aliquots of the three outlier healthy samples described above (Figure 4A, “PC2-high healthies”). As expected, the post-spin samples had much lower PC2 scores than the unspun samples (Figure 4A, PC scores), and much less total cfDNA in the >7,500bp bin (Figure 4A “Frac DNA”, 4-9% for post-spin samples and 38-48% for pre-spin samples). While they were both characterized by long fragments, the PC1-high and PC2-high groups had very different fragment length distributions (top of Figure 4A shows PC loadings). Likewise, while the long fragments in PC2-high samples had very low levels of CC-initiating motifs, the PC1-high samples had significant levels (Figure 4B), suggesting that the “hypofragmented” DNA in these cancers undergoes significant fragmentation in blood by DNASE1L3 after it is released from cells. Importantly, this analysis cannot resolve whether the hypofragmented DNA was released in circulation or *ex vivo* in the blood tube before extracting plasma.

PC3 was most enriched in fragments of length 75-145 bp and 245-295 bp (corresponding to short mono-nucleosomes and short di-nucleosomes, respectively) and consisted of a group of 17 “hyperfragmented” cancers (Figure 4A, “PC3-high cancers”). Longer fragments, including those greater than 1kb, were under-represented in these samples. Similar hyperfragmented cancers have been described previously both with Illumina sequencing [16,17] and ONT sequencing [24,25].

The remaining healthy and cancer samples had low values of all three PCs and were termed “healthy-like cancers” (Figure 4A). As described in earlier work, healthy-like cancers had significantly higher CC-initiating end motifs than the PC3-high cancers (Figure 4C). One intriguing aspect of the PCA analysis was that both the hypofragmented (PC1-high) and hyperfragmented (PC3-high) groups had elevated levels of cfDNA and H3.1 (Figure 4A, “DNA conc.” and “H3.1 nuc conc”) relative to healthy-like cancers, even if they had very low tumor faction (Figure 4A, “CAN ILLUMINA”).

The PCA analysis described above was run including the three “unspun” samples. To ensure that these unusual samples were not influencing the clustering of the cancers, we performed PCA with these samples excluded, which produced nearly identical versions of PC1 and PC3 (as the first two components), and identical grouping of all cancer samples (Supplementary Figure 4D-G).

### Hypofragmented DNA can result from cellular release during prolonged blood processing

Given that longer fragments can be the result of blood cell DNA release during processing, we sequenced plasma from a cohort of 89 pre-treatment neuroendocrine cancer patients that were varied in a key pre-analytic step. While all patients were from the same study with consistent patient characteristics, about 1/3 were from the “home hospital” close to the blood processing lab, while the remaining 2/3 were from one of several “external hospitals” in remote cities. Blood from the home hospital remained at room temperature for a maximum of 6 hours before processing, while blood from the external samples remained at room temperature for up to 24 hours during transport (Figure 5A).

**Figure 5:**
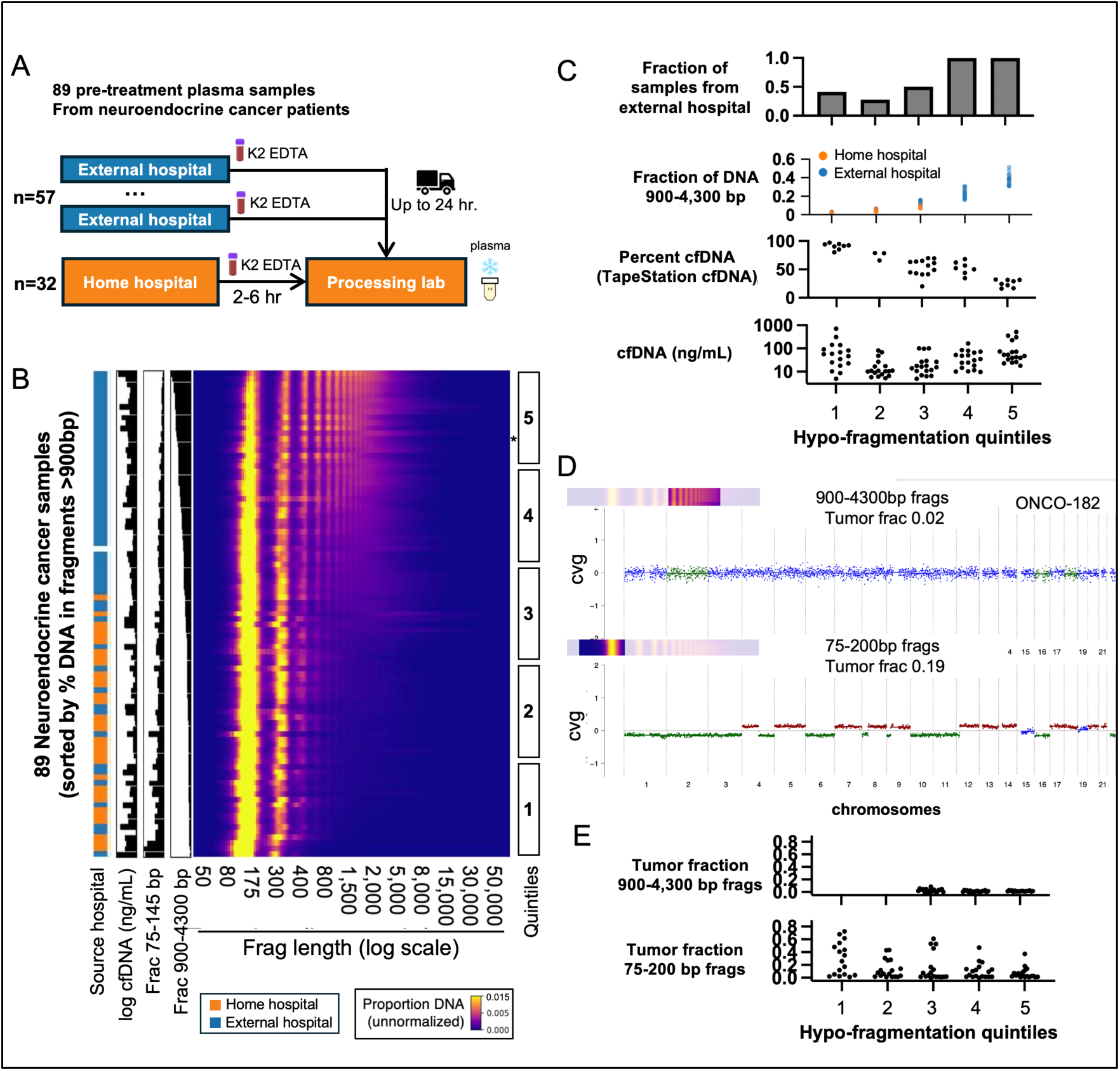
Hypofragmented DNA can be the result of cellular release during prolonged blood processing. (A) Neuroendocrine cancer blood samples collected either at the “home hospital” or several “external hospitals” were processed identically except for transport times. (B) Nanopore WGS of neuroendocrine samples, ordered by the fraction of DNA contained in fragments of 900-4300 bp. “log cfDNA” shows the log-transformed total plasma concentration of DNA, and “Frac 75-145 bp” and “Frac 900-4300 bp” show the fraction of sequenced DNA at in these two fragment length intervals. Five quintile bins are shown, with quintile #1 being the most hyperfragmented and bin #5 being the most hypofragmented. A representative hypofragmented sample (ONCO-182) is marked with an asterisk. (C) Comparisons of several features in samples in different hypofragmentation bins. (D) ichorCNA copy number plots for hypofragmented sample ONCO-182. The top plot is based on analysis including only 900-4300 bp fragments, and the bottom with only 75-200 bp fragments. (E) Tumor fractions for all samples calculated using 900-4,300 bp fragments (top) and 75-200 bp fragments (bottom). For 900-4,300 bp version, only the final 3 quintiles are shown because others have too few fragments to perform ichorCNA analysis.

We used ONT to sequence each sample to a median genomic depth of 1-2x (Supplementary Figure 5A) and analyzed fragment length distribution. When samples were sorted based on the percentage of DNA contained in long fragments of 900-4,300 bp, it was clear that many of the “external hospital” samples were enriched for these long fragments, whereas “home hospital” samples were not (Figure 5B). Furthermore, the length distribution was quite similar to many of the “hypofragmented” samples identified in our earlier pan-cancer cohort. Based on this sample ordering, we defined five equal bins with bin #1 having the lowest fraction of long fragments and bin #5 having the most (Figure 5B, right). Bins #4-5 were entirely from external hospitals and had 20-50% of DNA contained in long fragments (Figure 5C, top two charts). We also used Agilent TapeStation to size the DNA from these samples, which showed the “percent cfDNA” quantification (percent of fragments <200 bp) was very strongly associated with the quintile bins (Figure 5C, middle and Supplementary Figure 5B). As we observed in our pan-cancer cohort, cfDNA levels were increased in both the most hypofragmented and hyperfragmented bins.

In order to further verify that longer fragments were derived from blood cells in hypofragmented samples, we performed ichorCNA analysis based on fragment length subsets (Figure 5D-E). ichorCNA plots of one hypofragmented sample (ONCO-182, marked with an asterisk in figure 5B and shown in detail in Figure 5D) indicated significant cancer DNA fraction in typical 75-200 bp reads but undetectable levels of cancer DNA in the longer reads. The absence of cancer DNA in longer fragments was evident in all hypofragmented samples from bins #3-5 (Figure 5E, top). The apparent decrease in cancer DNA fraction even in shorter fragments in bin #5 relative to other bins (Supplementary Figure 5C) suggests that some contaminating DNA which begins as *ex vivo* long fragments may eventually be fragmented down to short fragments and dilute cancer fraction (consistent with in vitro experiments in blood serum, [36]).

### Hyperfragmentation is linked to reduced DNASE1L3-associated end motifs and elevated circulating cfDNA levels in the pan-cancer cohort

Hyperfragmentation occurred in 9 of the 12 cancer types in our pan-cancer cohort (Figure 4A) and a number of samples in our neuroendocrine cohort (Figure 5B). While this phenomenon has been documented in cancer [15–17,25], the process that underlies it remains poorly understood. To provide additional insights we performed feature clustering on the pan-cancer cohort based on the hyperfragmentation component (PC3). We first excluded the 7 PC1-high hypofragmented cancers, leaving 17 PC3-high cancers and 13 PC3-low cancers (Figure 6A). We calculated pairwise correlations between PC3 and six other tumor features and used these to perform hierarchical clustering, which defined three major feature clusters, shown in green, blue, and red (Figure 6A). We also mathematically split the total H3.1 and cfDNA levels into their tumor and non-tumor components based on ichorCNA tumor fraction (Figure 6B).

**Figure 6:**
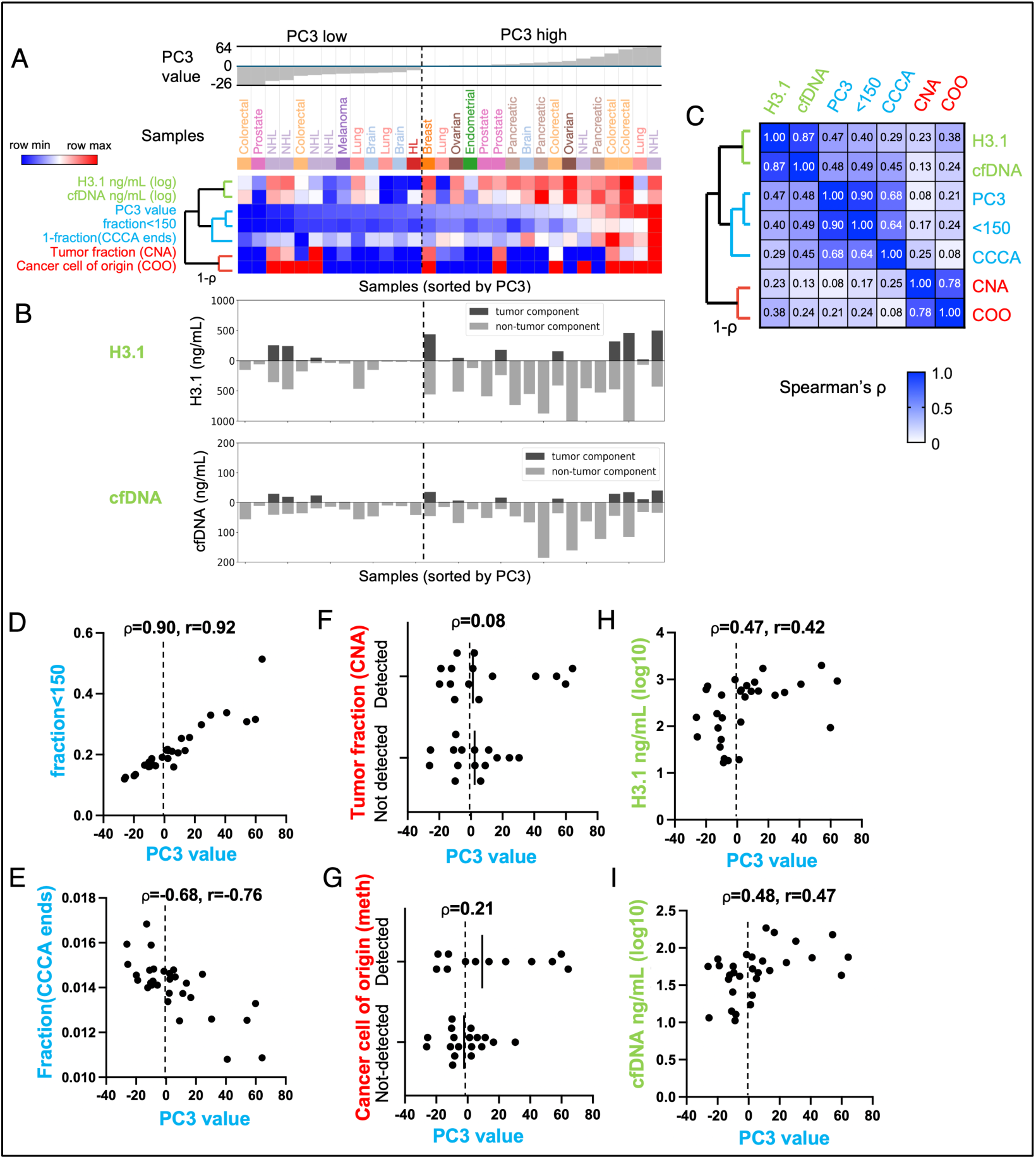
Hyperfragmentation is linked to reduced DNASE1L3-associated end motifs and elevated circulating cfDNA levels in the pan-cancer cohort. (A) 30 cancer samples from the PC3-high and PC1-3 low groups, ordered by PC3 value. Heatmap rows show sample features clustered by hierarchical clustering based on 1-ρ (Spearman’s rho). Heatmap rows are normalized to row min and max values. (B) Total H3.1 and cfDNA concentrations, mathematically split into the tumor and non-tumor component by multiplying by the ichorCNA tumor fraction to get the tumor component, and subtracting this from the total to get the non-tumor component. (C) Spearman correlation values for all pairwise comparisons. (D-I) PC3 values plotted against the following features: (D) fraction of short (<150 bp) fragments, (E) fraction of fragment ends with CCCA motif, (F) positive detection by ichorCNA, (G) positive detection by cancer methylation cell of origin, (H) H3.1 nucleosomes, and (I) total cfDNA.

The PC3 hyperfragmentation feature itself clustered into a central blue cluster, together with the fraction of fragments <150 bp, which is a commonly used definition for hyperfragmentation [17] (pairwise Spearman ⍴ of 0.90, Figure 6C). Also within this blue cluster was the DNASE1L3-associated CCCA end motif with an average Spearman ⍴ of 0.66 to the other two blue cluster features (Figure 6C and Figures 6D-E). The clustering of hyperfragmentation features together with the CCCA end motif is consistent with earlier findings that both hyperfragmentation and reduced CCCA frequency occur with reduced DNASE1L3 activity [13,19,35,37].

The two metrics that directly detected cancer DNA (copy number alteration, CNA, and methylation cell of origin, COO) were tightly clustered together with a Spearman ⍴ of 0.78 (Figure 6C, red cluster). While there were positive associations between these features and the hyperfragmentation (blue) cluster features, the average pairwise correlation between features in these two groups was only from 0.17 (Figure 6F-G) to 0.29 (Supplementary Figure 6A). Notably, these two clusters did not form a supercluster.

H3.1 nucleosome and cfDNA concentration were tightly clustered together with a Spearman ⍴ of 0.87 (Figure 6C, green cluster). These features formed a supercluster with the PC3/DNASE1L3 (blue) cluster (average Spearman ⍴ 0.43). While cancers with low PC3 hyperfragmentation values had somewhat variable cfDNA and nucleosome concentrations, those with high PC3 values almost always had elevated concentrations (Figure 6H-I). Interestingly, the cfDNA or H3.1 component attributable to non-cancer (i.e. blood) cells was highly elevated in almost all of these samples, whether or not the cancer component was (Figure 6B). This suggests a potential link between DNASE1L3-associated hyperfragmentation and the abnormally high levels of immune-derived cfDNA observed in cancer [4], and is consistent with similar results from a new study that suggests this is the result of a cancer inflammatory response [38].

Earlier studies observed an enrichment of cancer DNA in shorter fragments in hyperfragmented samples [16,17]. To measure this, we calculated CNA tumor fraction and methylation cell of origin directly in the shorter fragments. Of the 10 PC3-high cancers with sufficient read coverage, only 2 had measurable enrichment for cancer DNA, and neither was more than 1.5-fold enriched (Supplementary Figure 6B-G).

### Hyperfragmentation is linked to elevated cfDNA levels in additional cohorts

Next, we sought to investigate hyperfragmentation in our neuroendocrine cancer cohort. Our initial analysis showed a significant number of samples with elevated cfDNA level in the most hyperfragmented bin #1 (Figure 5B-C). To investigate this further, we performed a PCA analysis using the same method used for the pan-cancer cohort, which produced a single principal component PC1 that ordered samples from the most hyperfragmented to the most hypofragmented (Supplementary Figure 8A) and was strongly correlated to both short fragments 75-145 bp and long fragments >900 bp (Supplementary Figure 8B-C). Consistent with our pan-cancer findings, both the hypofragmented and hyperfragmented samples had consistently elevated cfDNA levels compared to samples with intermediate PC1 values (Figure 7A). While there was a strong correlation with tumor fraction in the hyperfragmented samples (Supplementary Figure 8A), the levels of cfDNA attributable to non-tumor origin were highly elevated irrespective of high levels attributable to tumor origin (Figure 7A).

**Figure 7:**
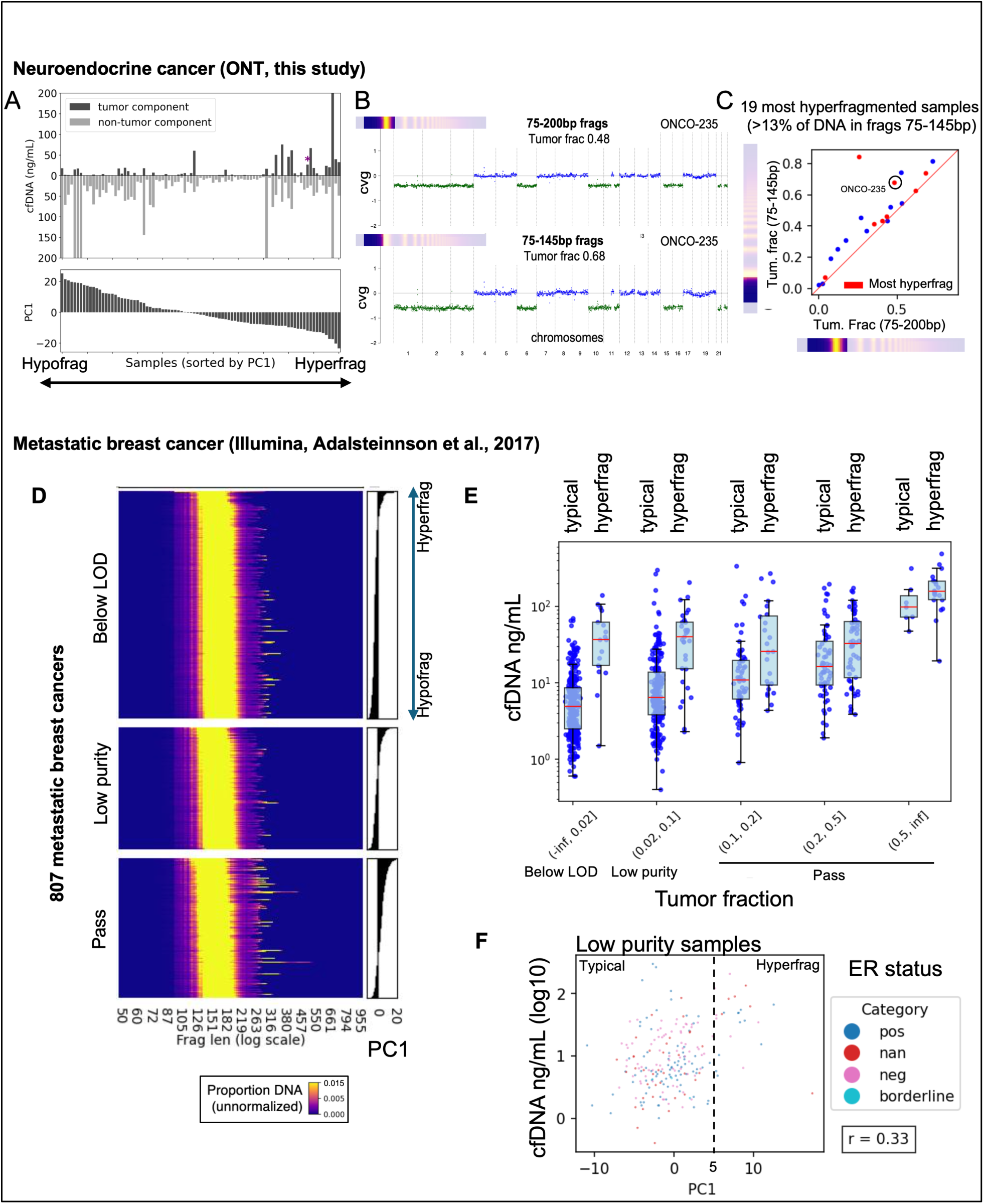
Hyperfragmentation is linked to elevated cfDNA levels in additional cohorts. (A) Principal Component Analysis was run on fragment lengths from the 89-sample neuroendocrine cohort, and the hyperfragmentation component (PC1) is shown in relation to total plasma cfDNA level. A representative hyperfragmented sample (ONCO-235) is circled. (B) ichorCNA is run separately on short fragments (75-145 bp) and mononucleosome fragments (75-200bp) on ONCO-235. (C) ichorCNA is repeated for all samples with >13% of DNA in 75-145 bp fragments, showing the tumor fraction in short vs. mononucleosome fragments. The 8 most hyperfragmented samples are shown in red. (D) Principal Component Analysis was run on 806 samples, and the first component (PC1) corresponded a fragment length spectrum with the most hyperfragmented samples having positive PC1 values, with samples grouped based on ichorCNA classifications defined in the earlier study (Figure 7). (E) Samples were divided into bins based on tumor fraction, and subdivided based on hyperfragmentation, (PC1<=5 being “typical” and PC1>5 “hyperfragmented”), and values of cfDNA concentration in plasma were plotted. (F) The bin corresponding to “Low purity” is shown in detail, colored by Estrogen Receptor status from the original publication.

In order to quantify the fraction of ultra-short cfDNA attributable to tumor origin in these samples, we inferred tumor fraction based on CNAs separately in 75-145 bp fragments vs. 75-200 bp fragments. A modest but significant enrichment of cancer DNA can be seen in some hyperfragmented samples, such as ONCO-235 (Figure 7B). However, in agreement with the results from our pan-cancer cohort, most of the hyperfragmented samples had only very slight enrichment if any for tumor-derived DNA, with only one sample having enrichment more than 1.5-fold (and this sample appears to have anomalous tumor fraction estimates, because the estimated ploidy strongly disagreed between the 75-145 bp and 75-200 runs). Our analysis suggests that tumor and blood cell DNA have similar levels of hyperfragmentation within an individual case, with only slightly higher fragmentation of tumor-derived DNA.

In order to validate the correlation between short fragments and elevated cfDNA levels in an independent study, we re-analyzed an Illumina-based WGS dataset consisting of 806 metastatic breast cancer cases [29]. We downloaded fragment length profiles and performed PCA as described above, which produced a single principal component PC1 that reflected hyperfragmentation (Figure 7D). We grouped samples into ichorCNA tumor fraction bins as described in the original paper - “Below LOD” (tumor fraction<0.02), “Low purity” (tumor fraction 0.02-0.1), and “Pass” (tumor fraction>0.1). Consistent with our neuroendocrine cohort, the high tumor fraction class (“Pass”) was strongly enriched for hyperfragmented samples (Figure 7D). We selected the hyperfragmented samples (PC1>=5) within each of the tumor fraction bins and plotted their cfDNA values vs. the “typical” samples (PC1<5). In all bins, hyperfragmented samples had higher cfDNA concentrations than typical samples, although this was most pronounced in the “Below LOD” and “Low Purity” bins. The “Low Purity” bin (shown in detail in Figure 7F) is especially important in this analysis because the “Below LOD” bin may include cancers that actually have high tumor fractions but lack any copy number alterations in their genomes. The Low Purity bin (0.02<tf<=0.1) and the next highest bin (0.1<tf<=0.2) both have readily detectable CNAs with low tumor fraction, and the hyperfragmented samples in these bins have strongly elevated cfDNA levels typically between 10-100 ng/mL, very similar to both the pan-cancer and neuroendocrine ONT cohorts.

## Discussion

In this study, we applied Oxford Nanopore (ONT) long-read sequencing to a diverse set of cancers to characterize circulating DNA and its fragmentomic patterns. We confirmed that ONT sequencing can classify cancer tissue of origin based on cell type–specific DNA methylation, as previously shown in a single cancer type [24], and extended this to multiple cancers. These findings support the potential of ONT for multi-cancer early detection (MCED) [6] and for determining tissue of origin in cancers of unknown primary (CUP) [39–41]. More importantly, our work provides a systematic survey of fragmentomic alterations in cancers with high cfDNA levels, leading us to identify four major classes of fragmentation patterns (Figure 8).

**Figure 8:**
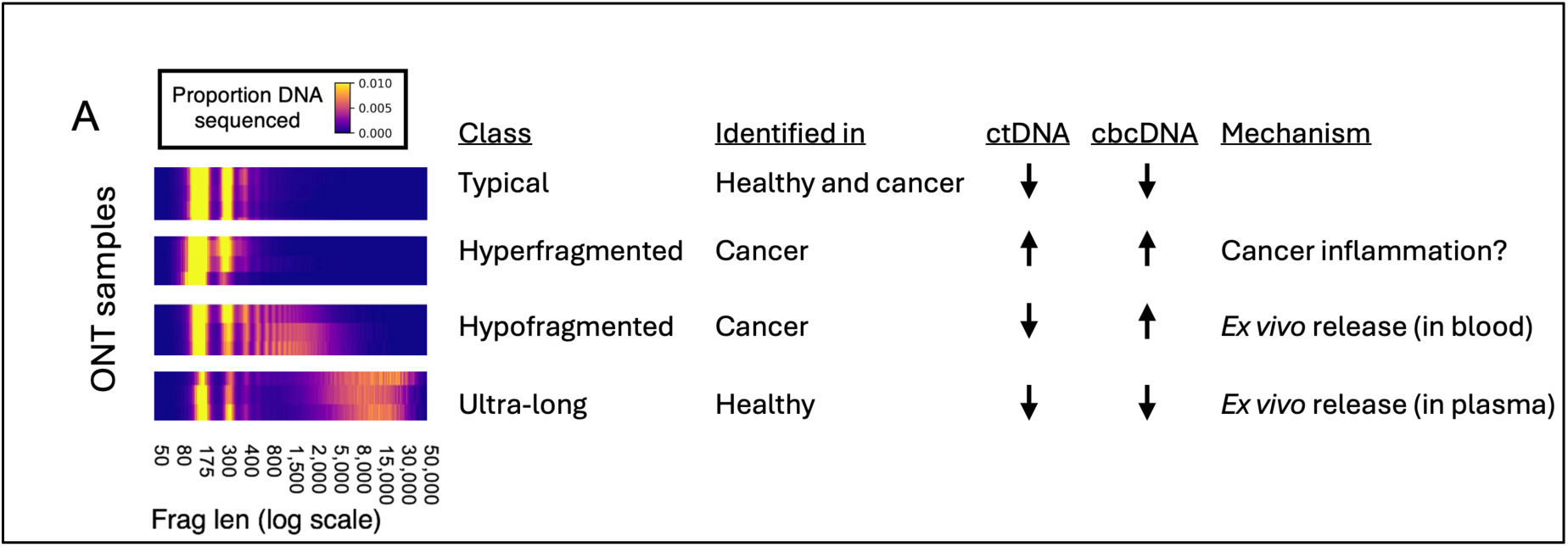
Four fragment length classes discovered in this study. ctDNA (circulating tumor DNA) and cbcDNA (circulating blood cell DNA) refer to the total concentration of DNA in the plasma sample.

We identified a distinct class of ultra-long cfDNA fragments (>7.5 kb) that were absent of DNASE1L3-associated end motifs and therefore unlikely to have been processed by circulating nucleases. These fragments were strongly reduced, though not entirely eliminated, by high-speed centrifugation and were only observed in one cohort. Their properties are consistent with ex vivo release from lysed blood cells during plasma thawing [33,34], representing a contamination class that can confound cfDNA analyses if not carefully controlled.

A second distinct class, which we term *hypofragmented*, was defined by excess fragments in the 1–4 kb range (5–25 nucleosomes). These fragments exhibited strong nucleosome phasing and retained DNASE1L3-associated end motifs. In our neuroendocrine cohort, hypofragmentation was strongly associated with prolonged blood storage at room temperature prior to plasma isolation. More than 40% of samples with storage times of 6–24 hours showed this signature, whereas samples processed rapidly did not. Because these long fragments lacked cancer-derived DNA, we conclude they are released from blood cells and cleaved by nucleases during blood processing.

These findings underscore the importance of pre-analytical handling in cfDNA studies, especially long-read sequencing where the bioinformatic filtering is possible but reduces sequencing coverage. Hypofragmentation can be partially detected by routine QC methods such as TapeStation analysis, and bioinformatic filtering can mitigate—but not fully eliminate—its impact. Notably, even shorter fragments from hypofragmented samples were diluted with ex vivo–derived cfDNA, suggesting contamination affects both long- and short-read sequencing approaches. Use of blood collection tubes containing fixatives (e.g., Streck tubes) may be preferable when immediate processing cannot be ensured.

Although most hypofragmented signatures identified here appeared to arise *ex vivo*, long cfDNA fragments have been associated in other contexts with biological processes. Prior studies have observed similar signatures in patients with inherited DNase1L3 mutations [13,19,27,35], Dnase1l3-deficient mice [19], pregnancy [26,28], and cancer [23,26]. Additionally, longer cfDNA fragments have been linked to high cfDNA concentrations in healthy plasma [42]. Importantly, the strong nucleosome phasing and DNASE1L3-associated end motifs in these fragments make them difficult to distinguish from in vivo sources such as neutrophil extracellular traps (NETs) [36], which have been implicated in cancer-associated inflammation and elevated cfDNA levels [43–45], and likely generate similar sized fragments [36,45]. These findings highlight the need to disentangle pre-analytical from biological origins when studying long cfDNA fragments.

Hyperfragmentation (<145 bp fragments) was present across both cancer cohorts and is consistent with previous reports using short- and long-read sequencing [15–17,20,25,30]. Our findings here are generally in agreement with earlier results indicating that hyperfragmentation is modestly to strongly correlated with higher tumor fraction [16,17,30]. Earlier work showed that hyperfragmentation can occur in inflammatory conditions such as systemic lupus erythematosus (SLE) [35,46], and a new study revealed nearly identical signatures in cancer and other inflammatory diseases [38]. In this study, the hyperfragmentation signature correlated with inflammatory blood protein markers, strongly implicating innate immune processes as a shared driver of hyperfragmentation and elevated cfDNA levels [38].

Reduced activity of DNASE1L3 may play a central role in this phenotype. Familial SLE caused by DNASE1L3 mutations is associated with cfDNA hyperfragmentation [35], and common DNASE1L3 missense variants have also been linked to similar signatures [13]. Together with our evidence here and other new evidence [38] that hyperfragmentation is correlated with high cfDNA concentrations, these findings suggest that altered stoichiometry between cfDNA and circulating endonucleases contributes to this phenotype. Alternatively, hyperfragmentation may reflect a shared cfDNA release mechanism across cancer and immune cells, potentially through phagocytosis as suggested by the high levels of innate immune markers such as MPO in these cases [38].

Previous studies have shown that cancer-derived cfDNA is more hyperfragmented than cfDNA from blood cells [16,17]. In our data, this effect was inconsistent: some patients showed modest differences between cancer- and blood-derived fragments, while others showed no difference, similar to patient data from [17]. By contrast, xenograft studies consistently demonstrated greater hyperfragmentation of tumor DNA [17,25]. A systematic mutation-based analysis found that fragment shortening was only a weak predictor of cancer origin across solid tumor types (AUROC 0.51–0.61) [47]. Given that the hyperfragmentation process is likely inflammatory and not cancer-specific, we propose that chromatin changes intrinsic to cancer cells may sensitize cancer DNA to cleavage [48–50], especially at specific long-range chromatin domains [18]. This could explain why hyperfragmentation is observed in some, but not all, cancer-derived fragments.

Our results reveal that ONT sequencing can capture a broad spectrum of cfDNA fragmentomic patterns, including artifact-derived and biologically relevant signatures. We identified ultra-long and hypofragmented fragments primarily as *ex vivo* artifacts, whereas hyperfragmentation likely reflects inflammatory processes affecting both cancer and immune cells. Together with ONT’s ability to profile DNA modifications and tissue of origin, these findings suggest that long-read sequencing can provide unique biological insights and broaden the scope of circulating DNA biomarkers, but proper pre-analytic standards will be required to take full advantage of this technology. Given its simple library preparation and potential for rapid, point-of-care deployment [51,52], ONT sequencing may emerge as a powerful tool for both research and clinical applications.

## Conflicts

BPB, SAE, CW, JC, JVT, MH, and TKK are present or past employees or contractors for VolitionRx and hold VolitionRx stock. VolitionRx holds intellectual property in the area of cancer liquid biopsy, including circulating DNA and Oxford Nanopore sequencing. LP, MP and TW are employees of University of Lyon 1 and Hospices Civils of Lyon have no conflict of interests.

## Author contributions

TKK and BPB conceived the study and co-supervised the research. SAE, CW, JC, and TKK processed samples from the pan-cancer cohort. MO, MP, LPG, and TW processed blood samples from the Neuroendocrine Tumor cohort, and MH facilitated transfer of frozen DNA. SAE, CW, and JC performed library construction and sequencing. BPB, JC, JVT, and performed bioinformatic analyses. BPB drafted the manuscript, with TKK, JC, and SAE contributing. All authors have reviewed the manuscript.

## Acknowledgements

The authors thank Daniel Halter and Jacques Avaux for support with cloud computing and Nextflow workflow infrastructure. We also thank Andrew Retter, Jake Micallef, Catherine Mallalieu, and Gaetan Michel for helpful comments.

## Data availability

We have deposited three levels of data. “Level 1”: BAM files before filtering steps were modified by replacing all single-nucleotide variants with the base in the reference genome hg38, and all insertion variants with Ns (see Methods). These “anonymized” BAM files allow all fragmentomic and copy number analysis to be performed without exposing any personally identifying genetic variants. Level 1 data is deposited in Short Read Archive BioProject ID PRJNA1106648. “Level 2”: In order to provide combined fragmentation and methylation information, Biscuit [53] was used to create “epiBED” files (see Methods). epiBED files contain each read on a separate line, with each DNA methylation call on the read. epiBED files can be used for combined methylation/fragmentomic analysis, and are compatible with the CelFiE-ISH software [32] that was used for cell of origin deconvolution. Level 3: DNA methylation BED files. BED files created by modkit (see Methods) provide one line for each CpG covered and can be used with bedtools and a number of other DNA methylation analysis tools. Level 2 and Level 3 data are deposited at Zenodo with DOI 10.5281/zenodo.11092490.

## Code availability

Most analysis was performed using publicly available tools, with all software version numbers and command line settings listed in the Methods section. For several in-house scripts that were developed to analyze fragmentation patterns, source code is deposited at Zenodo with DOI 10.5281/zenodo.11092490.

## Methods

### The pan-cancer and healthy cohort, plasma collection and processing

Plasma samples were procured through the following Commercial Biobanks: Disovery Life Sciences (DLS), Innovative Research, and Bay Biosciences. All donor and biospecimen information is available in the Supplementary Data File 1. Selection of 52 self-reported healthy individuals for Illumina sequencing were selected based on donor age criteria: an even distribution of ages from 40 to 90 years old. Selection of 207 cancer cases was based on Stage III/IV status and diversity of cancer types. Selection of healthy samples was based on volume of plasma available and the age of the donor. Samples were evenly distributed across ages with 10 samples in each of the following age groups: 40-49, 50-59, and 60-69 years old. 15 samples were procured in the age group 70-79, and 5 samples in the age group 80-89. Healthy samples were only considered on a lack of cancer diagnosis. Cancer samples were selected based on late stage diagnosis (III or IV), type of cancer, treatment status (untreated) and large sample volume.

All three commercial biobanks collect blood in EDTA tubes, then isolate the plasma by centrifuging at 1300-2000 xg for 10-15 minutes. The plasma is then frozen at −80°C. Exceptions to this protocol are listed in Supplementrary Data File 1. Plasma samples were received and subsequently stored at −80°C. Aliquots were thawed at room temperature (RT) for up to 2 hours, depending on the aliquot volume. Plasma was then spun at 14000 x *g* for 2 minutes at room temperature and transferred to a new tube avoiding any pelleted fraction.

### Neuroendocrine tumor cohort plasma collection and processing

The cohort included 89 patients from the “MGMT-NET” trial (Walter et al. 2024). Patients were age 18 or older with a histologically proven well-differentiated neuroendocrine tumor (NET) from the pancreas or lung, grades 1-3, metastatic or locally advanced, and unresectable. Blood samples were taken prior to any treatment. Blood was collected and processed as described previously [54]. Briefly, 3 × 10 mL of whole blood was collected in K2 EDTA tubes.

32 “Home hospital” blood samples were collected at the Edouard Herriot Hospital, Hospices Civils de Lyon, France, and sent by pneumatic transport to the East hospital laboratory before being transferred by road through at room temperature via an internal shuttle to our team at the South hospital laboratory for pre-analytical processing. These samples were processed within 6 hours after blood collection. 57 “External hospital” blood samples were collected using the same protocol at various other participating hospitals in France, and transferred by road at room temperature via an internal shuttle to the South hospital laboratory. These samples were processed within 24 hours after blood collection.

All blood samples were centrifuged for 10 min at 1,600 g, and the cell pellet was discarded. The supernatant was then centrifuged at 6,000 g for 10 min, and the resulting plasma was stored in 2-mL cryotubes and kept at −80°C until thawing.

### Nucleosome Quantification

Nucleosome levels in plasma were quantified using the Nu.Q^®^ H3.1 assay developed by Volition for use on the IDS i10 instrument, using 250 uL of plasma according to the manufacturer’s recommendations. Samples were completed either in duplicate or triplicate. Raw RLU (Relative Light Units) were converted to concentrations of nucleosomes (ng/mL) using the provided kit standards, and replicate measurements were averaged.

### DNA Extraction and Basic Characterization

#### Pan-cancer and healthy cohort

All DNA extractions were completed with the QIAamp^®^ Circulating Nucleic Acid Kit (Qiagen, catalogue # 55114) or with the QIAamp^®^ MinElute ccfDNA Kit (Qiagen, catalogue # 55204). When possible, the extraction was completed with 1 mL plasma (if less, 1X PBS was used to bring volume to 1 mL). Additional extractions (if DNA levels were not sufficient for ONT) were carried out with up to 5 mL plasma, according to the manufacturer’s protocol. The Qiagen protocol was followed per the manufacturer’s recommendations with the following adjustments: samples were eluted in 30 µL heated (60°C) elution buffer and the elution step was also completed at 60°C to increase DNA yield. Samples were quantified with Qubit and total DNA levels were calculated based on plasma volume input. DNA profiles were confirmed using BioAnalyzer or Tapestation (Agilent).

#### Neuroendocrine cohort

1-101.5-100 ng of DNA used as input for Nanopore sequencing were extracted using the QIAamp^®^ Circulating Nucleic Acid Kit (Qiagen, catalogue # 55114) according to manufacturer’s protocol. Samples were quantified with the Qubit 2.0 Fluorometer and the Qubit dsDNA HS Assay Kit (Life Technologies, Q32854). When necessary, if DNA levels were not sufficient for ONT, additional extractions were carried out on back-up samples of plasma. DNA fragment profiles were evaluated using the Agilent 2100 BioAnalyzer (Agilent Technologies) with the DNA HS kit (Agilent Technologies, 5067-4626 & 5067-4627) and confirmed with Tapestation (Agilent) before ONT sequencing, either with the cfDNA tape kit or the genomic tape kit.

### Illumina Sequencing and mapping

When possible, DNA from the basic characterization step was used for Illumina Sequencing. If more was needed, additional DNA from 1-5 mL of plasma was extracted using the QIAamp^®^ Circulating Nucleic Acid Kit (Qiagen, catalogue # 55114). Illumina sequencing libraries were constructed using single-stranded DNA SRSLY® PicoPlus DNA NGS Library Preparation Kit (ClaretBio – Cat: CBS-K250B). Libraries were sequenced by Discovery Life Sciences using the NovaSeq 6000 S4 200 cycle flow cell. Genome mapping to hg38 was performed using the Illumina BaseSpace DRAGEN. Mapping quality and filtering metrics are available as Supplementary Data File 1.

### Oxford Nanopore (ONT) Sequencing, pan-cancer cohort

#### Pan-cancer and healthy cohort

Samples were prepped for Nanopore sequencing using the 109 library prep kit and the 104 and 114 barcoding kits (ONT catalogue: SQK- LSK109, EXP-NBD104, and EXP-NBD114, respectively). The Oxford Nanopore Technology (ONT) protocol was used, with the following adjustments: In the DNA repair and end-prep section, DNA input was decreased to 30 ng, and incubation time was increased from 5 min at 20C and 5 minutes at 65C to 30 minutes at each temperature. Throughout the protocol, the AMPure bead: reaction ratio was changed from 1:1 to 3:1, and ethanol wash volumes were increased from 200 uL to 400 uL. When sequenced, additional adaptations were made to the input and amount of library loaded per flow cell as needed based on the success of the previous sequencing runs and the number of pores available on the flow cell. R9.4.1 flow cells were sequenced on the MinION Mk1c, with sequencing runs lasting for 16-72 hours, depending on the health of the flow cell. Samples were sequenced individually but were still barcoded to avoid possible contamination. Plasma input and DNA input are included in Supplementrary Data File 1.

#### Neuroendocrine cohort

Sequencing was performed as with the pan-cancer and healthy cohort, with the following changes. For library construction, we used the Oxford Nanopore Technologies Native Barcoding Kit v. 14 (SQK-NBD114-96). For sequencing, we used PromethION R10.4.1 flow cells on the P2 Solo sequencer.

### ONT basecalling, mapping, and DNA modification calling

#### Pan-cancer and healthy cohort

Barcode demultiplexing and quality filtering were performed by MinKnow using FAST model. “Passed” fast5 files were re-processed using Dorado v. 0.2.4+3fc2b0f using the SUP basecalling model dna_r9.4.1_e8_sup@v3.3 and the Remora modification calling model dna_r9.4.1_e8_sup@v3.3_5mCG@v0, which resulted in unaligned BAM files including DNA modification tags. These were mapped to the UCSC hg38.analysisSet assembly (https://hgdownload.soe.ucsc.edu/goldenPath/hg38/bigZips/analysisSet/) using Minimap v. 2.24-r1122 with “minimap2 -y -t 2 -ax map-ont” and filtered with samtools v. 1.14 with “samtools view -e ([qs] >= 10) && (mapq >= 20)”. Duplicates were marked with samtools v.1.17 “samtools markdup”, and supplementary and secondary mappings were removed using “samtools view -F 0xD04”.

For global methylation analysis and CpG Island methylation analysis, methylation BED files were created using modkit v. 0.1.5 with the command “modkit pileup --cpg -- combine-strands --ignore h --filter-threshold 0.9 --bedgraph”. For methylation deconvolution, Minimap BAM files were used directly (details below). BED files are available in the Zenodo repository listed in Data Availability.

#### Neuroendocrine cohort

Nanopore sequence data as processed as above for the pan-cancer and healthy cohort, with the following changes. Base and modification calling, demultiplexing, and QC were performed using the ONT Dorado basecaller v. 0.4.1 with the dna_r10.4.1_e8.2_400bps_hac@v4.2.0_5mCG_5hmCG modification calling model. Alignment was performed with minimap2 version 2.24-r1122 as above. DNA modifications were extracted using Oxford Nanopore modkit 0.1.13 with the same flags as above.

### Copy number alteration analysis

#### Default settings

Analysis of CNAs was performed using ichorCNA v. 0.5.0. Default parameters were used with all chromosomes 1-22, with a genomic bin size of 1E6. The following other parameters were used:

centromere=’GRCh38.GCA_000001405.2_centromere_acen.txt’, scStates=’’,repTimeWig=’’,ploidy=2,maxCN=7,includeHOMD=FALSE,estimateNo rmal=TRUE,estimatePloidy=TRUE,estimateScPrevalence=TRUE,altFracThresho ld=0.05,genomeBuild="hg38",genomeStyle="UCSC",flankLength=100000,txnE=0.9999999,chrs="c(’chr1’,…,’chr22’)",chrTrain="c(’chr1’,…,’chr22’)",normal="c(0.5)"

CNA was considered “detected” if tumor fraction was estimated to be greater than 0.

#### Sensitive settings

Sensitive settings were used for the pan-cancer cohort (Illumina and ONT) as an alternative parameter set (Supplementary Figure 2). For the neuroendocrine cohort, sensitive settings were used as the default analysis. Sensitive parameters had the following differences from the default parameters:

maxCN=3,estimateScPrevalence=FALSE,
normal="c(0.2,0.3,0.4,0.5,0.6,0.7,0.8,0.9,0.95,0.97,0.98,0.99)"

#### Panel of Normals, pan-cancer healthy cohort

A panel of normals (ichorCNA parameter “normal_panel”) was used for the pan-cancer cohort as an alternative setting for both Illumina and ONT (Supplementary Figure 2). For the Illumina cases, this used the full set of 52 healthy samples, sequenced using the same single-stranded library protocol as the cancer cases. For the ONT cases, this used the full set of 21 healthy samples.

#### Panel of Normals, neuroendocrine cohort

A panel of normals (ichorCNA parameter “normal_panel”) was used for the neuroendocrine cohort as the default setting. In this case, we did not have a matched set of healthy samples, so we used a subset of tumor samples that got tumor fraction of 0 with ichorCNA default settings without a Panel of Normals. We clustered all samples based on normalized coverage in 1 Mb bins and confirmed that these samples formed a tight cluster separate from the samples that got non-zero tumor fractions without the panel of normals. This resulted in a set of 47 (of the 93 total) samples which were used as the panel of normals. In analyses of specific fragment length bins (Figure 6G-J), we used a panel of normal created using only fragments of the specified size (we found that these could be quite different from the panel of normals created using all fragments).

### Fragment length analysis

Fragment lengths were extracted from ONT Minimap BAMs using the script “fragmentationReports.py”, which uses Pysam and defines the fragment length based on the primary alignment, calculating the difference between the start and end coordinates on the reference genome. Fragment length distributions for PCA analysis were created by defining bins as 10^x where x contains a range 50 to 50,000. Raw fragment lengths are log transformed and assigned to the closest bin by rounding to the nearest increment of 10^0.01. The proportion of fragments is defined as the number of fragments in a given bin divided by the total number of fragments in all bins. For the proportion of total DNA, each fragment in a bin is multiplied by the number of base pairs in the fragment and summed to get the bin base pair total. Each bin total is then divided by the sum of all base pair sums in all bins. The code to perform this analysis is available as the script “fraglens_to_histogram.py”.

In order to perform CNA, end motif analysis, and methylation cell of origin analysis in specific fragment length bins, Minimap BAMs were filtered using the script “stratifyBamByFraglen.py”, which uses the same Pysam code as above to calculate the fragment length of each read, and write that read to the output BAM file only if it is within the correct length range.

### End motif analysis

ONT Minimap BAMs were processed using the script “fragmentationReports.py”, which uses Pysam to perform the following analysis. We collect all reads that contain a perfect match to the final 5 base pairs of the ONT adapter sequence (“CACCT”) as the final sequence in the soft-clipped portion of the read. We then collect the first 5 base pairs of the aligned portion of the read, which is by definition adjacent to the adapter sequence. This is the “end motif”. If any of the adjacent 5 base pairs have a basecalling quality less than 20, the end is not counted. We count each of the two ends of each fragment as independent observations.

### Principal Component Analysis (PCA) of fragment length distributions

DNA proportions were calculated for fragment length bins as described above. PCA was performed using the sklearn.decomposition Python package. For main figures 2-5, PCA was performed on column normalized versions of the fragment length distributions.

These were defined by taking the proportion of DNA in each bin for of a given sample and calculating a z-score based on the mean and standard deviation of the bin across all 21 healthy samples (not including the 3 “unspun” healthy samples). PCA in the main figures included 3 “unspun” healthy samples. PCA in Supplementary Figure 4D-G excluded these samples.

### Cancer methylation signatures

For the “global” hypomethylation signature, we used the mean methylation value at 495,252 genomic CpGs across the genome which have been associated with rapid methylation loss during replication in cancer [55]. These were taken from the file zhou_NN_scores_PMD_top_bottom_10ptile.0based.hg19.bed.gz [56] and lifted over to hg38. For the CpG island signature, we used the mean of all CpGs contained within 3,329 CpG islands that were significantly hypermethylated within human cancer data from The Cancer Genome Atlas (TCGA) project [57].

### Methylation Cell of Origin (COO) analysis

We used a version of CelFiE-ISH package v0.0.2 (https://gitlab.com/methylgrammarlab/deconvolution_models),[32] modified to take Minimap BAM files as input. We used the command:

deconvolution --model uxm --min_length 2 --u_threshold 0.25 --modbam_qual 0.9 --percent_u U_atlas.hg38.tsv -m input.bam -i U250.hg38.tsv -b -- cpg_coordinates hg38.analysisSet.CGmotifs.modkit.merged.0based.bed.gz -- epiformat modbam --outfile output.txt

The “--modbam_qual 0.9” setting was used to filter out any modification base with a modification probability score less than 0.9. We used the 31 cell types from the WGBS atlas (Loyfer et al., 2023), as described in the CelFiE-ISH paper [32]. We summed individual cell types to create composite cell type groups such as “Gastrointesintal-Ep” (’Gastric-Ep’+’Colon-Ep’+’Small-Int-Ep’). All composite cell type groups are listed in Supplementary Figure 2. For Figure 2, each cell type was normalized by calculating the Z-score using the mean and standard deviation across all healthy and cancer samples.

### Creation of anonymized BAM files

We used script bamReplaceSeqByRef.py with “-ref hg38.analysisSet.fa”, the same UCSC reference genome used for alignment. Files are available in the Zenodo repository listed in Data Availability, and code is available in the Zenodo repository listed in Code Availability.

### Creation of Biscuit EpiBED files

Biscuit [53] v.1.4.1-dev was used with the command “biscuit epiread -M -b 0 -m 0 -a 0 -5 0 -3 0 -y 0.9 -L 1000000 hg38.analysisSet.fa”, where the reference is the same UCSC reference genome used for alignment. Files are available in the Zenodo repository listed in Data Availability.

### Statistical tests

Statistical tests, jitter plots, and scatterplots were generated with GraphPad Prism v. 10.2.1.

## Supplementary Tables

Supplementary Table 1: Metadata, cfDNA and H3.1 nucleosome quantification, and Illumina WGS ichorCNA features for the full screening dataset.

Supplementary Table 2: Metadata, cfDNA and H3.1 nucleosome quantification, and top-level sequencing features for the ONT R9.4.1 pan-cancer cohort

Supplementary Table 3: Detailed fragmentomic features for the ONT R9.4.1 pan-cancer cohort

Supplementary Table 4: Detailed DNA methylation features for the ONT R9.4.1 pan-cancer cohort

Supplementary Table 5: Metadata, cfDNA quantification, and top-level sequencing features for the ONT R10.4.1 Neuroendocrine cancer cohort

Supplementary Table 6: Detailed fragmentomic features for the ONT R10.4.1 Neuroendocrine cancer cohort

## Supplementary Figures

**Supplementary Figure 1:**
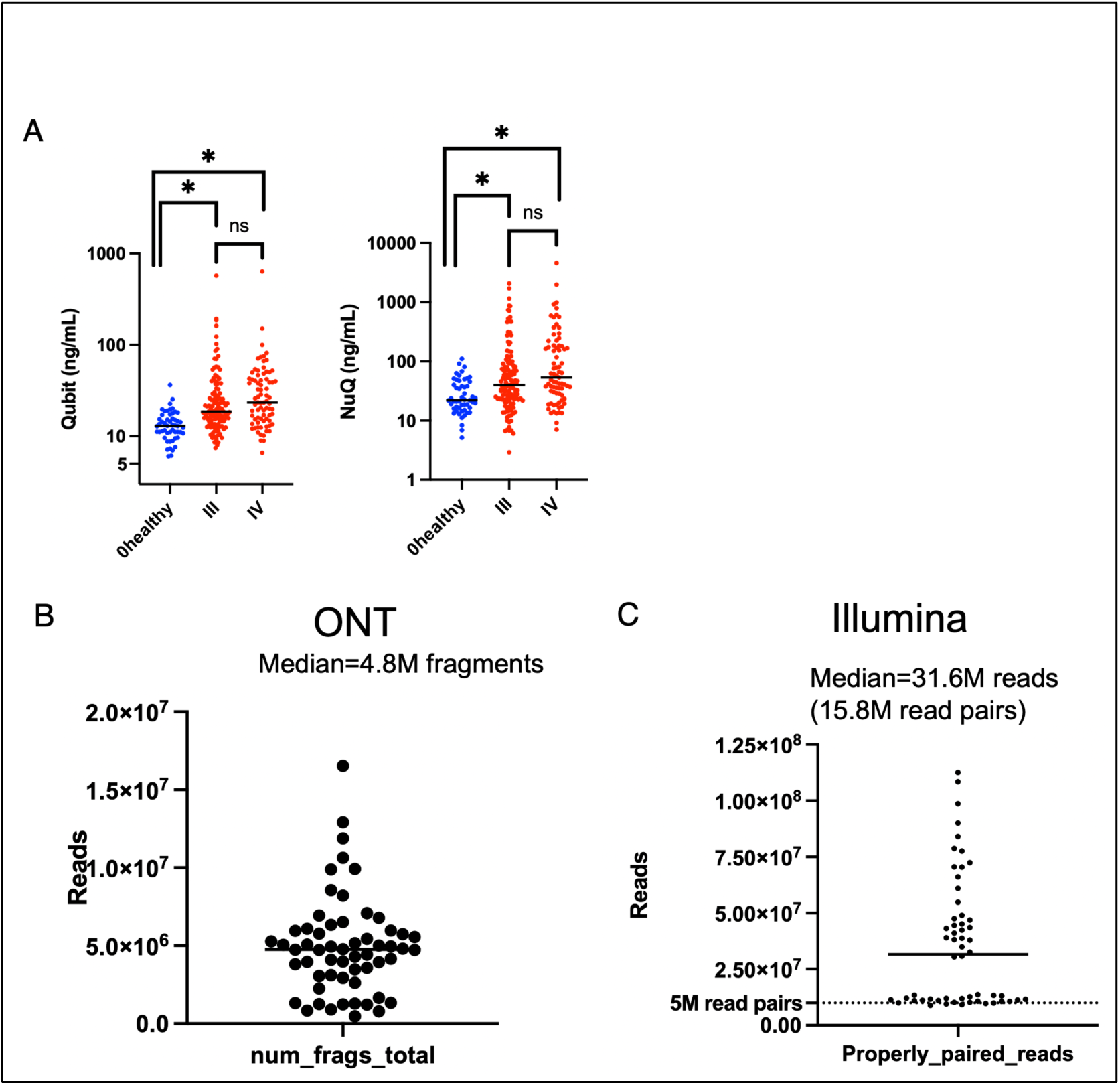
(A) cfDNA levels (left) and H3.1 nucleosome levels (right) in the screening dataset of 236 cancer samples and 52 healthy samples. (B) Read counts for 58 cancer and healthy samples from the pan-cancer healthy cohort, sequenced using ONT MinION Mk1c. Each fragment consists of a single read. (C) Read counts for the matched 58 cancer and healthy samples sequenced using Illumina NovaSeq. Each fragment consists of a single read pair. Panels A-B were performed using a two-tailed, unpaired t-test. Panel A p-values 0.015 (healthy vs. Stage III) and 0.011 (healthy vs. Stage IV), panel B p-values 0.011 (healthy vs. Stage III) and 0.016 (healthy vs. Stage IV).

**Supplementary Figure 2:**
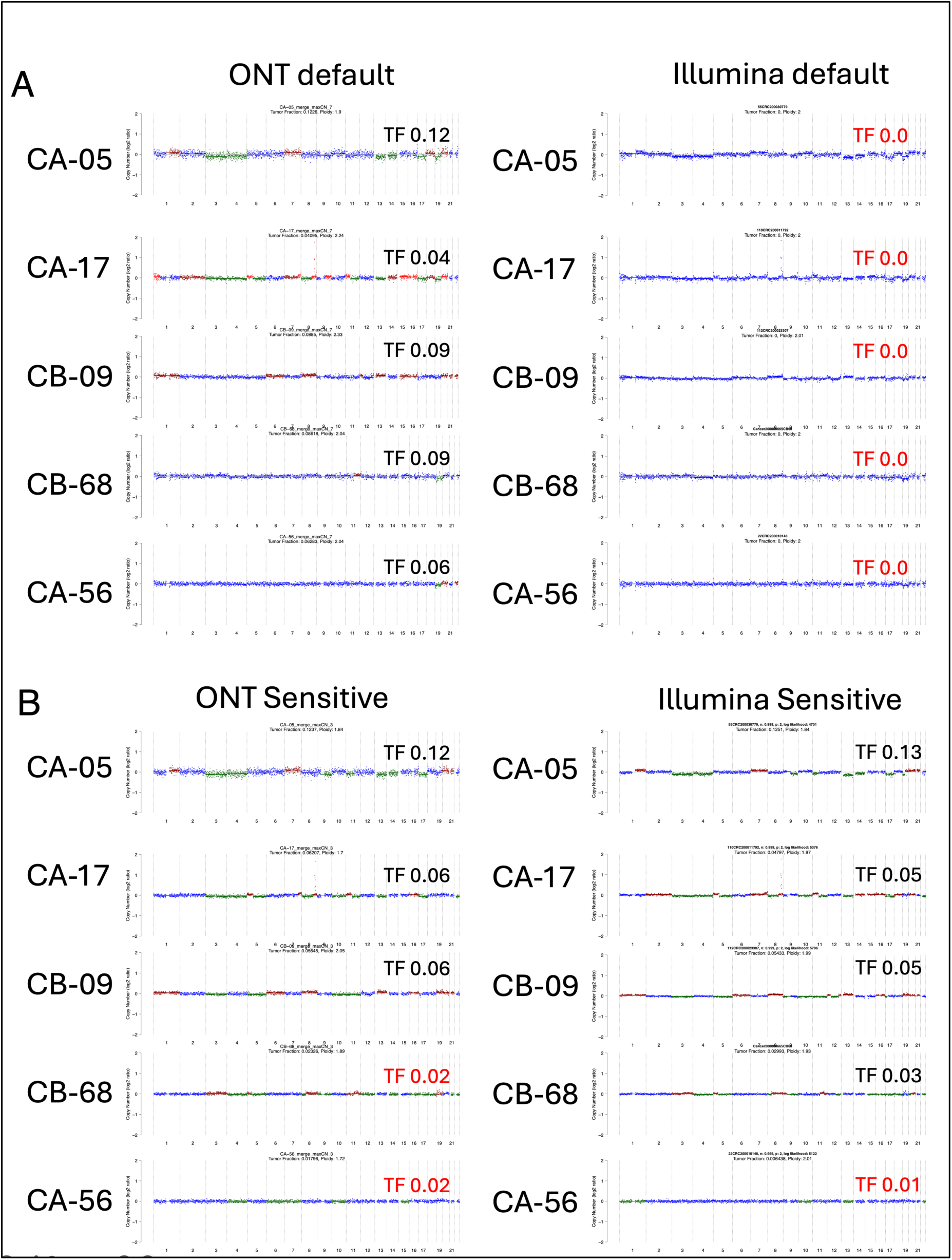
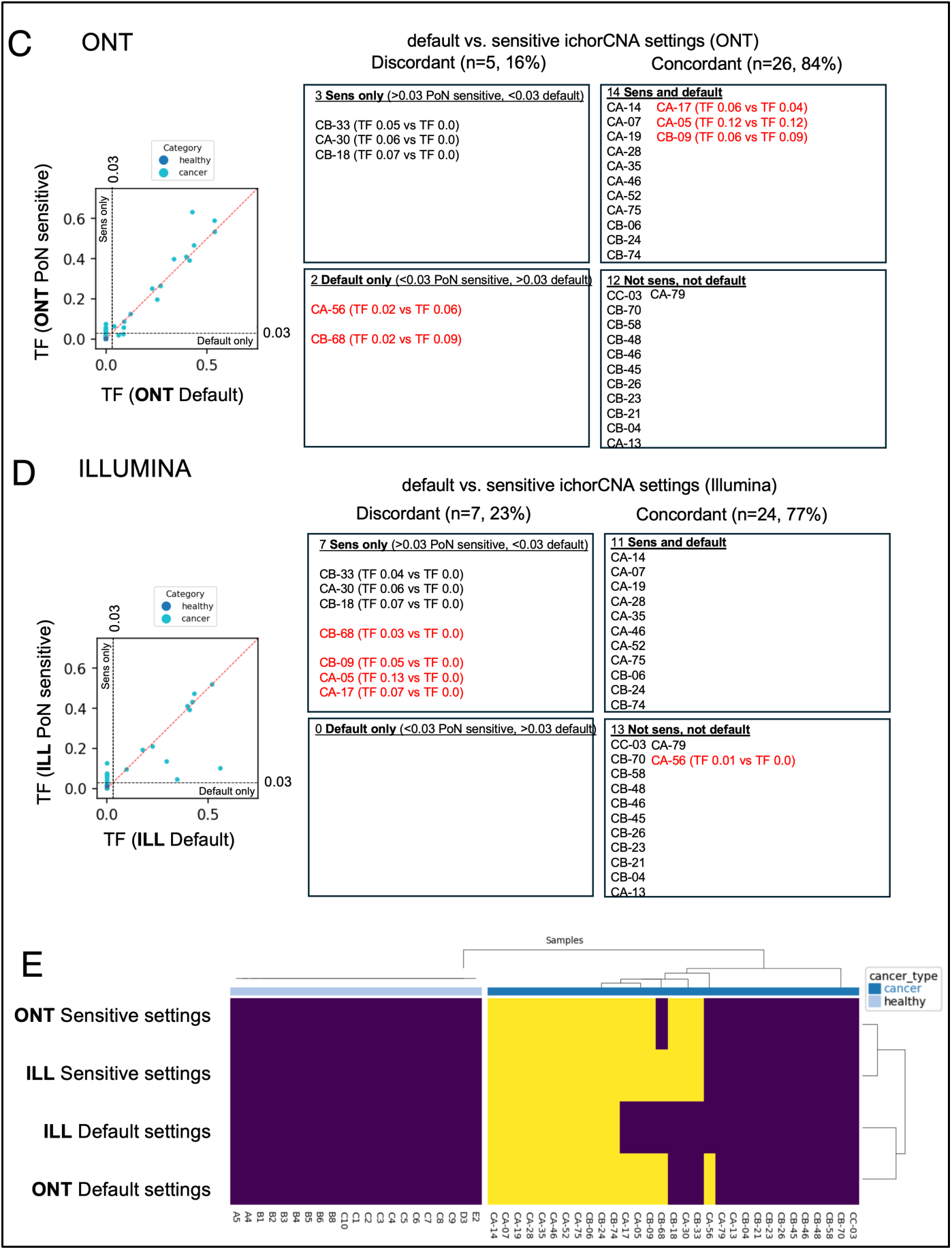
ichorCNA default settings vs. sensitive settings. (A) ichorCNA copy number plots for five samples that were only called by ONT and not by Illumina using ichorCNA default settings (“ONT-only” cases). (B) Same five samples, with ichorCNA sensitive settings. “TF” refers to estimated tumor fraction. (C) Differences in ONT samples between default and sensitive settings, for the 31 cancer samples run on both platforms. Scatter plot on left shows estimated tumor fractions, with dotted lines drawn at 0.03, the recommended limit of detection for ichorCNA. Boxes on the right are based on binarizing each sample as detected vs. not detected in each condition and dividing samples into boxes based on those samples that are discordant between default and sensitive settings (5 samples) and those that are concordant (26 samples). Cases listed in red are those that change boxes in the Illumina data below. (D) Same as C, but based on the Illumina samples. Red samples are those that are assigned to different boxes in the ONT analysis in C. (E) is a summary of C and D, where blue boxes have tumor fraction below the level of detection, and yellow boxes are above the level of detection.

**Supplementary Figure 3:**
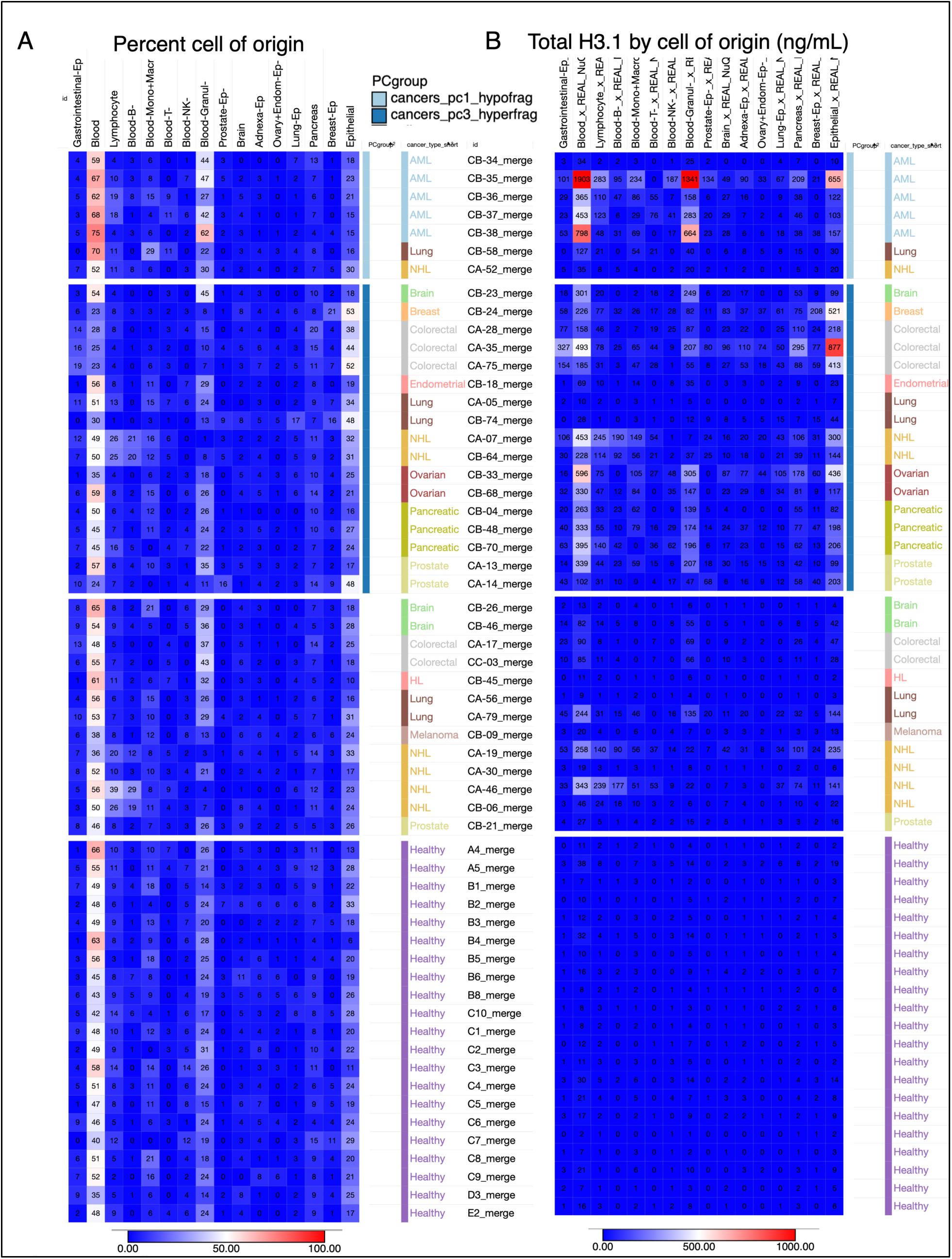
Methylation-based cell of origin (COO). (A) For each sample, the non-negative least squares (NNLS) method of CelFiE-ISH was used to estimate the percentage of the mixture derived from each of 31 individual cell types. We then combine several cell type groups by summing percentages, with cell type groups as follows: Gastrointestinal-Ep: ’Gastric-Ep’,’Colon-Ep’,’Small-Int-Ep’ Blood: ’Blood-T’,’Blood-NK’,’Blood-Mono+Macro’,’Blood-Granul’,’Blood-B’,’Eryth-prog’ Lymphocyte: ’Blood-T’,’Blood-NK’,’Blood-B’ Adnexa-Ep: ’Ovary+Endom-Ep’,’Fallopian-Ep’ Lung-Ep: ’Lung-Ep-Bron’,’Lung-Ep-Alveo’ Pancreas: ’Pancreas-Duct’,’Pancreas-Acinar’,’Pancreas-Delta’,’Pancreas-Beta’,’Pancreas-Alpha’ Breast-Ep: ’Breast-Basal-Ep’,’Breast-Luminal-Ep’ Epithelial: ’Bladder-Ep’, ’Breast-Basal-Ep’, ’Breast-Luminal-Ep’, ’Colon-Ep’, ’Fallopian-Ep’, ’Gastric-Ep’, ’Head-Neck-Ep’, ’Kidney-Ep’, ’Lung-Ep-Alveo’, ’Lung-Ep-Bron’, ’Ovary+Endom-Ep’, ’Prostate-Ep’, ’Small-Int-Ep’, ’Thyroid-Ep’ Since these groups are not mutually exclusive, the percentages do not sum to 100%, although the 31 primary cell type percentages do. For instance, Blood-B is shown as an individual cell type but also contained in both “Lymphocyte” and “Blood”. (B) Each cell type or cell type group percentage is multiplied by the total concentration of H3.1 nucleosomes (ng/mL) for the sample. The color scale is trimmed at a maximum value of 1,000 ng/mL.

**Supplementary Figure 4:**
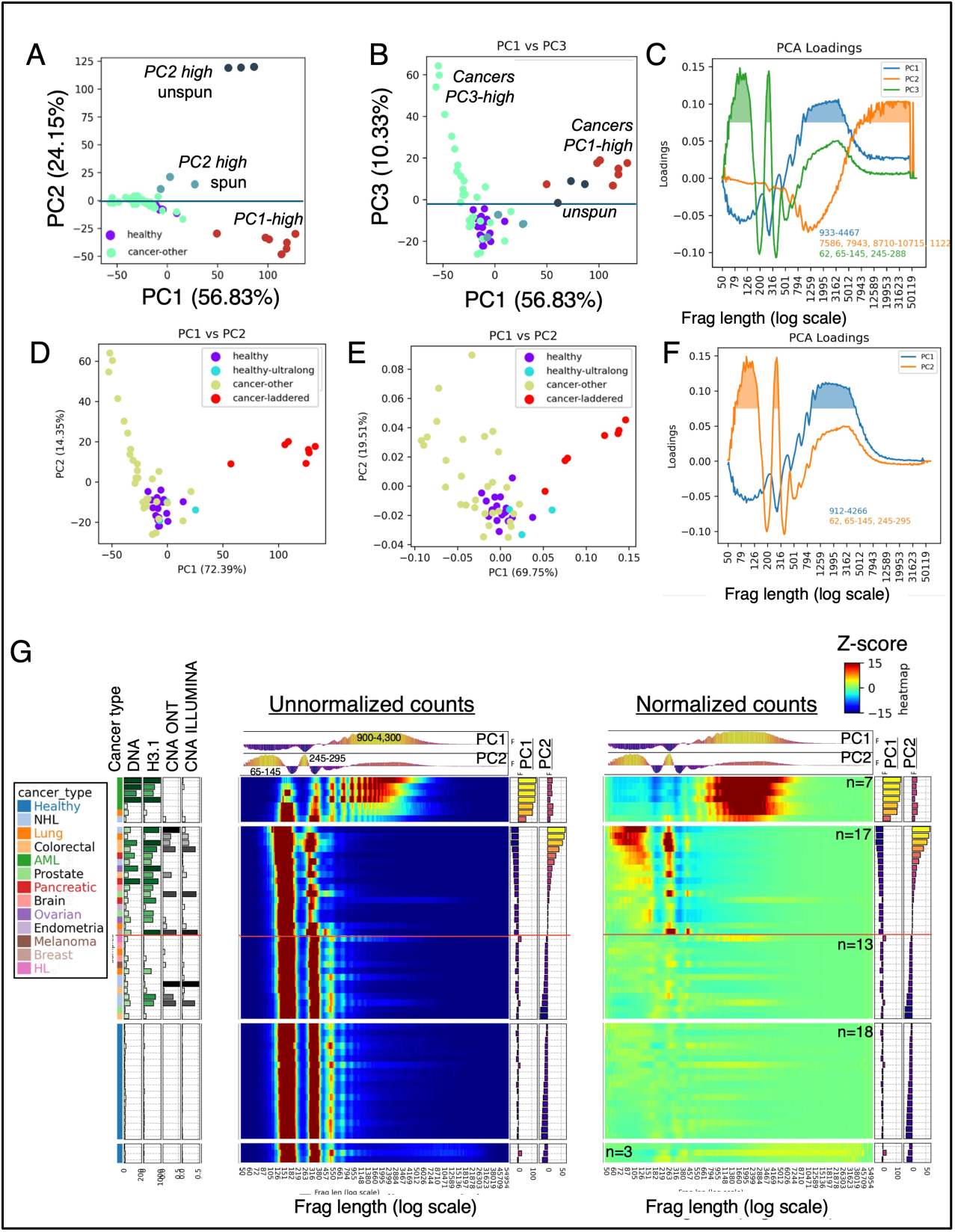
Principal Component Analysis of fragment length profiles. (A-C) correspond to the PCA presented in the main Figure 4. For this analysis, 58 healthy and cancer samples were included, plus the “unspun” samples of the 3 healthy volunteers with ultra-long fragments (E2, C10, D3). The input vector for each sample included one value for each fragment length bin, which represented the fraction of DNA in that bin, z-score normalized by the mean and standard deviation of the fractions of all healthy samples for the same bin (not including the “unspun” samples). (A-B) PC1, PC2, and PC3 values for each sample. In (B) a line is drawn between the PC3-high hyperfragmented cancer group and the normal-like cancer group in the main Figure 4. (C) shows the PCA loadings for each bin for each of the top 3 PCs. An arbitrary cutoff of 0.075 was used to define the bin ranges strongly associated with each PC, which are listed inside the figure. (D-G) An alternative PCA where the 3 unspun samples were omitted. The PC1 vs. PC2 plot in (D) is nearly identical to the PC1 vs. PC3 plot in the (B) panel. (E) An alternative PCA where the 3 unspun samples were omitted, and raw percent of DNA values were used rather than z-score normalized versions. (F) PCA loadings, defining the bin ranges strongly associated with each of the top 2 PCs, using the same cutoff as the (C) panel. (G) Sample ordering based on the alternative PCA.

**Supplementary Figure 5:**
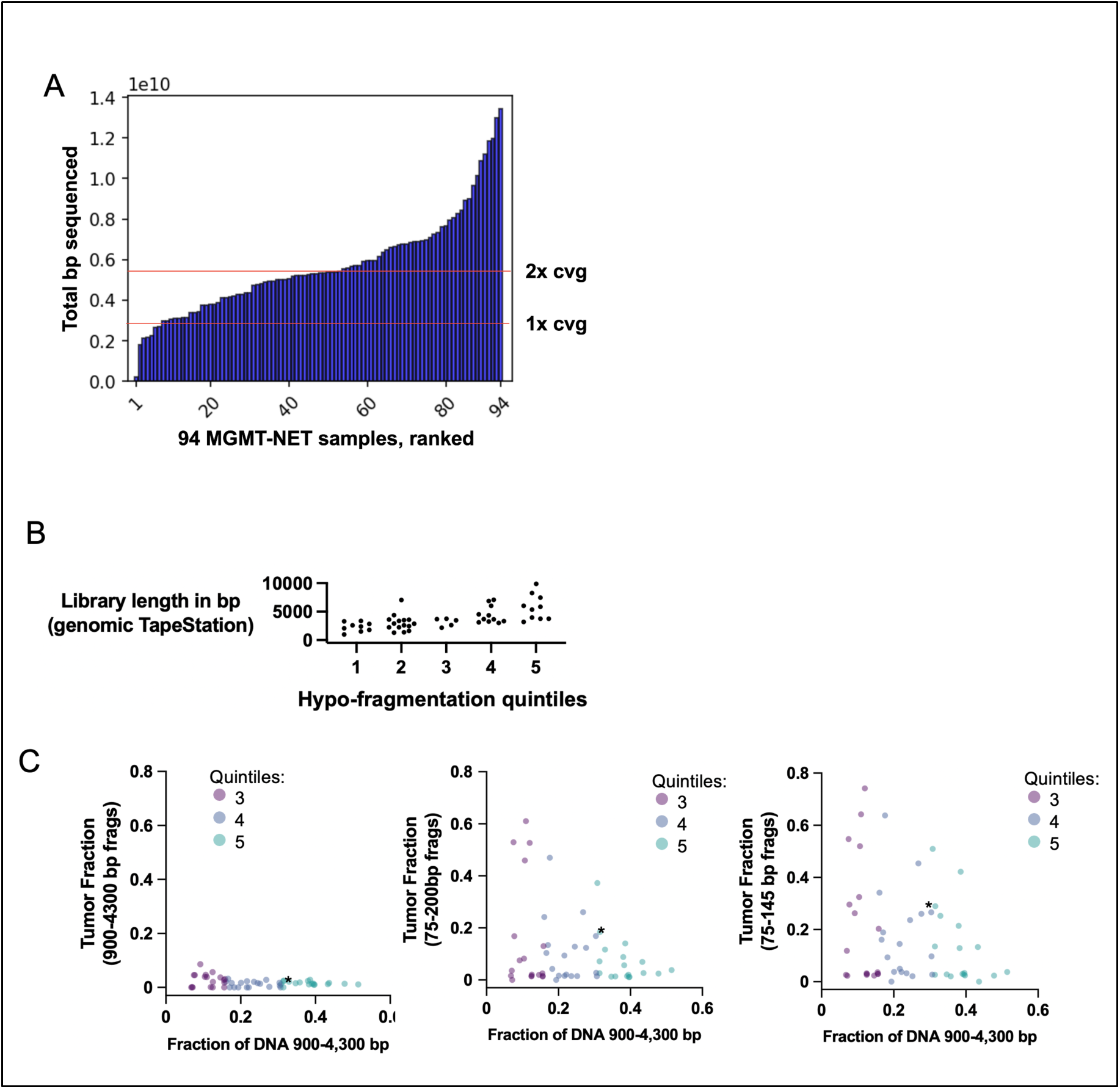
ONT neuroendocrine cohort. (A) 94 sequenced samples, ordered by the total number of base pairs in all sequenced reads. 5 samples were not included in subsequent analysis. (B) Samples where pre-sequence DNA sizing was performed using the “genomic” tape for the Agilent TapeStation system. Quintiles are those defined in Figure 5, from the least hypofragmented (quintile 1) to the most (quintile 5). (C) Hypofragmentation (Fraction of DNA in fragments 900-4,300bp) is plotted against ichorCNA tumor fraction estimates. In each case, the ichorCNA analysis is performed with a specific set of reads (900-4300bp fragments on the left, 75-200bp fragments center, and 75-145 bp fragments right). Coloring is based on the same hypo-fragmentation quintiles from Figure 5.

**Supplementary Figure 6:**
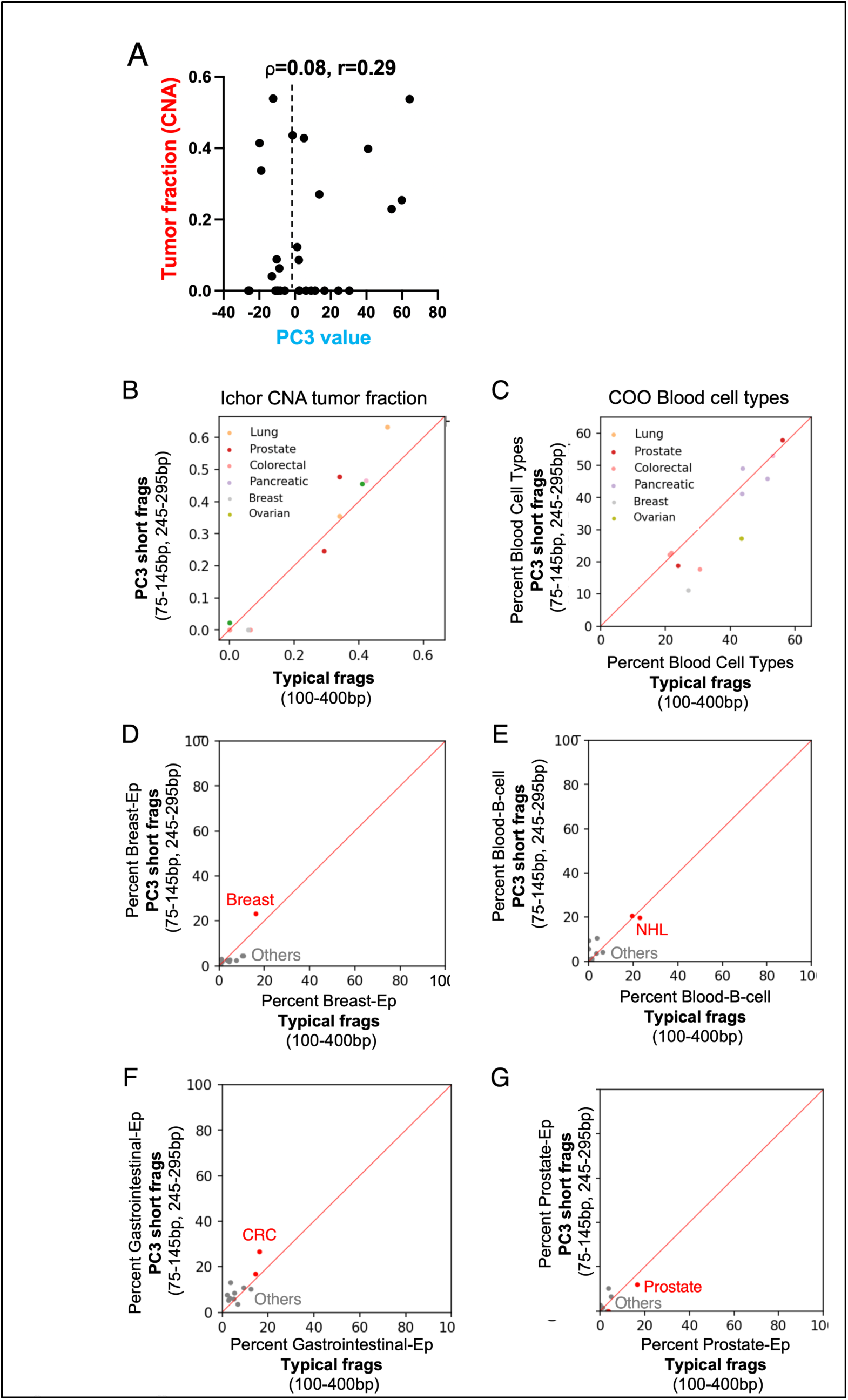
Associations between hyperfragmentation and cancer origin. (A) Alternative version of the binarized data shown in Figure 5E. Each point is one of 30 cancer samples (the 7 PC1-high hypofragmented cancer samples are omitted). The y axis is the tumor fraction estimated from ichorCNA based on the ONT sequence data. (B-G) Cancer cell fraction in shortened fragments in hyperfragmented samples. Of the 17 PC3-high samples (PC3>-2), 7 had less than 1 million fragments in either of the PC3-associated length ranges 75-145 or 245-295. We used the remaining 10 to perform ichorCNA and methylation-based cell of origin (COO) analysis. (A) ichorCNA was run using only typical fragments (100-400bp) and fragments in the PC3-associated length ranges. (B-F) Methylation cell of origin was computed for combined Blood cell types (B) as well as cell types which represented the correct cell of origin: Breast-Ep for Breast cancer (C), B-cell for Non-Hodgkins Lymphoma (D), Gastrointestinal-Ep for Colorectal cancer (E), and Prostate-Ep for Prostate Cancer (F)

**Supplementary Figure 7:**
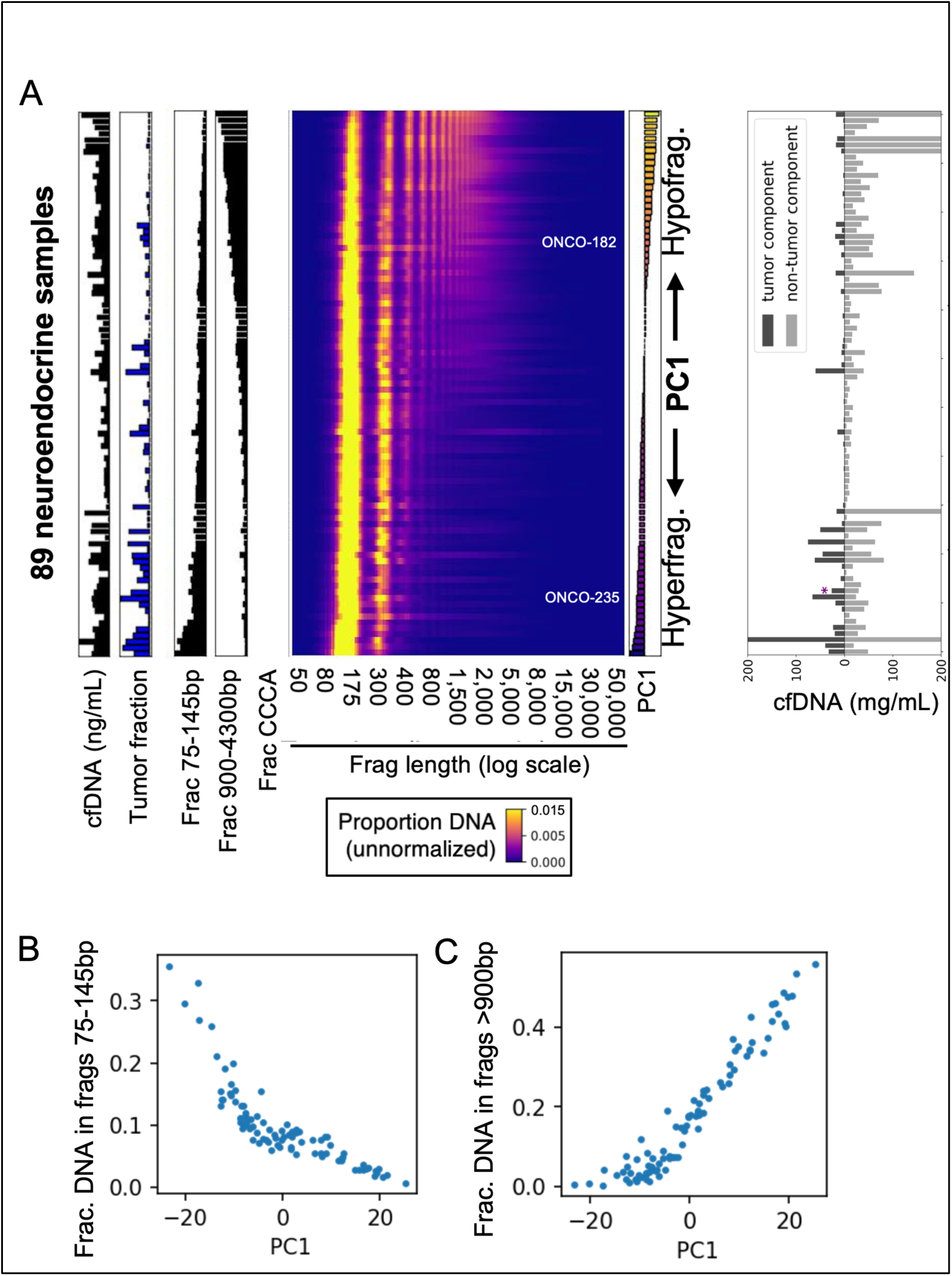
Principal Component Analysis of neuroendocrine samples based on fragment length. Samples are ordered by the first component (PC1), which separates hyperfragmented samples (bottom) from hypofragmented samples (top). (B-C) PC1 value plotted against the fraction of DNA in short fragments (B) vs. long fragments (C).

